# Macroevolutionary inference of complex modes of chromosomal speciation in a cosmopolitan plant lineage

**DOI:** 10.1101/2023.09.05.556433

**Authors:** Carrie M. Tribble, José Ignacio Márquez-Corro, Michael R. May, Andrew L. Hipp, Marcial Escudero, Rosana Zenil-Ferguson

## Abstract

- The effects of single chromosome number change—dysploidy—mediating diversification remain poorly understood. Dysploidy modifies recombination rates, linkage, or reproductive isolation, especially for one-fifth of all eukaryote lineages with holocentric chromosomes. Dysploidy effects on diversification have not been estimated because modeling chromosome numbers linked to diversification with heterogeneity along phylogenies is quantitatively challenging.
- We propose a new state-dependent diversification model of chromosome evolution that links diversification rates to dysploidy rates considering heterogeneity and differentiates between anagenetic and cladogenetic changes. We apply this model to *Carex* (Cyperaceae), a cosmopolitan flowering plant clade with holocentric chromosomes.
- We recover two distinct modes of chromosomal evolution and speciation in *Carex*. In one diversification mode, dysploidy occurs frequently and drives faster diversification rates. In the other mode, dysploidy is rare and diversification is driven by hidden, unmeasured factors. When we use a model that excludes hidden states, we mistakenly infer a strong, uniform positive effect of dysploidy on diversification, showing that standard models may lead to confident but incorrect conclusions about diversification.
- This study demonstrates that dysploidy can have a significant role in large plant clade speciation despite the presence of other unmeasured factors that simultaneously affect diversification.

## Introduction

Unveiling the primary drivers of diversification remains one of the most important goals in evolutionary biology (Sauquet & Magallón 2018). Hundreds of studies have focused on estimating changes in plant diversification processes through time (Magallón & Castillo 2009), across clades (Magallón et al. 2019), or in association with trait evolution (Helmstetter et al. 2023). Chromosome number changes and rearrangements are particularly likely to influence lineage diversification (Freyman & Höhna 2018). Yet, most plant diversification studies that address chromosome evolution focus only on the role of polyploidy, rather than on dysploidy—gains and losses of single chromosomes—which does not involve large DNA content changes (*i*.*e*., dysploidy, gains or losses of single chromosomes; Escudero et al. 2014, Mandáková & Lysak 2018). A recent review of trait-dependent diversification in angiosperms, for example, cites seven studies linking speciation and polyploidy (Helmstetter et al. 2023), only one which considered dysploidy linked to diversification (Freyman & Höhna 2018). While dysploidy has the potential to influence lineage diversification through effects on recombination or reproductive isolation, the macroevolutionary effects of dysploidy on speciation and extinction remain unknown, and far fewer macroevolutionary studies have focused on speciation and extinction as a result of dysploidy.

### Dysploidy and holocentric chromosomes

There are two classic models of chromosomal speciation in a macroevolutionary context. In the hybrid-dysfunction model, dysploidy is linked to speciation events. Under this model, dysploidy causes an immediate reproductive barrier, as reproduction between individuals with different chromosome numbers would cause problems during meiosis. Thus, most—if not all—dysploidy events across a phylogeny would occur cladogenetically. Alternatively, the recombination-suppression model posits that chromosomal rearrangements may become fixed in lineages via either drift or selection (as some rearrangements may physically link adaptive loci or locally reduce recombination). Under this model, dysploidy would evolve primarily anagenetically since fission and fusions do not necessarily restrict gene flow; rather, they evolve within populations and lineages (Baker & Bickham 1986). The two above-described models are not mutually exclusive. In a recent review, Lucek et al. (2022) introduce a third option—the hybrid-dysfunction/recombination-suppression model, under which dysploidy evolves both anagenetically and cladogenetically (see Fig. 4 in Lucek et al. 2022). While populations may be able to continue interbreeding despite some dysploidy events (which then may become fixed in the lineage), other dysploidy events may cause speciation, either because of an accumulation of differences that eventually leads to incompatibility or because of the genomic signature of the dysploidy event itself. We describe additional theory on dysploidy and macroevolution in Supplemental Section S1.

Holocentricity—having chromosomes without clear centromeres/primary constrictions—is distributed broadly across the Tree of Life, including 18 different lineages in animals, plants, and rhizaria (Escudero, Márquez-Corro & Hipp 2016, Márquez-Corro, Martín-Bravo, Pedrosa-Harand, Hipp, Luceño & Escudero 2019), approximately 15–20% of eukaryotic species (Márquez-Corro et al. 2018). Two particularly diverse holocentric clades show extraordinary chromosome number variation: the insect order Lepidoptera (de Vos et al. 2020) and the angiosperm sedge family Cyperaceae (2*n* = 4−224; Márquez-Corro et al. 2021, Márquez-Corro, Martín-Bravo, Spalink, Luceño & Escudero 2019). Holocentric chromosomes, instead of having kinetochore activity concentrated in a single point (*i*.*e*., at a single centromere in monocentric chromosomes), have centromeric regions distributed along the whole chromosome where the kinetochores assemble in most of the organisms with such chromosome type (Márquez-Corro, Martín-Bravo, Pedrosa-Harand, Hipp, Luceño & Escudero 2019, Marques et al. 2015). In monocentric chromosomes, many chromosome fissions are expected to result in a loss of genetic material during meiosis and inviable gametes, as chromosome fragments without centromeres are unable to segregate normally, and fusions are generally inherited by combining two telocentric chromosomes (Robertsonian translocations; Robertson 1916). Holocentric chromosomes, by contrast, allow chromosome fragments to segregate normally during meiosis (Faulkner 1972). Holocentricity may promote chromosome number variation via fission and fusion, as these changes are expected to be neutral or nearly so in holocentric organisms (Márquez-Corro, Martín-Bravo, Pedrosa-Harand, Hipp, Luceño & Escudero 2019). Thus, holocentric organisms provide a unique system in which to study chromosomal speciation (Lucek et al. 2022). Nonetheless, of the few applied macroevolutionary studies that address the role of dysploidy in lineage diversification, most have focused on monocentric chromosomes, which have a single centromere (*e*.*g*., Ayala & Coluzzi (2005), Freyman & Höhna (2018), but see relevant work on Lepidoptera de Vos et al. (2020), among others).

*Carex*—the largest genus in Cyperaceae—is particularly well suited to studying the effect of dysploidy on plant diversification because all *Carex* have holocentric chromosomes, the genus represents 40% of the third most species-rich monocot family (among the tenth in angiosperms, POWO 2023), and well-developed phylogenetic and chromosome number datasets for the genus facilitate macroevolutionary studies (Martín-Bravo et al. 2019, Márquez-Corro et al. 2021). In *Carex*, karyotype evolves mainly through fusion, fission, and translocations, in contrast to other sedge lineages where karyotype evolves through both dysploidy and polyploidy (Márquez-Corro, Martín-Bravo, Spalink, Luceño & Escudero 2019, Elliott et al. 2022, Shafir et al. 2023). *Carex* also has exceptional variability in chromosome number, ranging from 2*n* = 10 to 2*n* = 132 (Márquez-Corro et al. 2021). *Carex* has experienced several rapid radiations (Martín-Bravo et al. 2019), and shifts in optimal chromosome number are thought to have played a role in some of these radiations (*e*.*g*., *Carex* sect. *Cyperoideae*; Márquez-Corro et al. 2021, Hipp 2007).

Previous studies that have tested for a correlation between dysploidy and diversification (*e*.*g*., Márquez-Corro et al. 2021) have relied on models that fail to account for alternative sources of variation in diversification rates—*e*.*g*., morphological traits, climatic niche, or biotic interactions—that are not the study’s focal trait (in our case, chromosome number; Beaulieu & O’Meara 2016). Models that fail to account for alternative sources of variation in diversification rates have high type-I error rates (Rabosky & Goldberg 2015) because they misattribute underlying diversification-rate variation caused by unmeasured factors to variation due to the states included in the model. In other words, the null hypothesis of those models assumes that there is no underlying diversification-rate variation. This high error rate has motivated the development of hidden-state diversification models that account for underlying diversification-rate variation that is not caused by the focal trait (Beaulieu & O’Meara 2016, Caetano et al. 2018). These models not only more accurately test for associations between the focal trait and diversification, but also provide an opportunity to model how focal evolutionary processes vary across the phylogeny. Here, we design a new model of joint chromosome evolution and lineage diversification that incorporates process variation in chromosome number evolution and disentangles the effects of dysploidy from other unobserved factors that may also affect diversification rates (described in detail below). We apply our model to the most recent *Carex* time-calibrated phylogeny with chromosome number information that contains over 700 taxa and more than 50 states (Martín-Bravo et al. 2019, Márquez-Corro et al. 2021).

We test for a detectable effect of chromosome gains and losses on the rate of species formation in a large, diverse group of plants with high variation in chromosome number: *Carex* (Cyperaceae, Poales). We demonstrate that dysploidy is sometimes associated with faster rates of lineage formation, but diversification and dysploidy play out through two evolutionary modes across *Carex*. In one mode, chromosome rearrangements evolve rapidly and are linked to new species formation. In the other mode, rearrangements evolve less frequently and are not associated with new species formation; instead, other factors—hidden states unmeasured in our analysis—likely drive the formation of new species. Our results demonstrate the complexity of the diversification process and the important role of genomic rearrangements in determining biodiversity patterns at geological timescales. Furthermore, we illustrate how computational and statistical advances in modeling permit increasingly nuanced models that better represent the underlying complexity of biological systems.

## Materials and Methods

### Modeling chromosome number evolution

We developed the Chromosome number and Hidden State-dependent Speciation and Extinction model (ChromoHiSSE) to test the interaction between single chromosome number change, speciation, extinction, and diversification heterogeneity. Our model can be considered a natural extension of the ChromoSSE model (Freyman & Höhna 2018). Under the ChromoSSE model, lineages evolve independently under a continuous-time Markov chain (CTMC) that describes changes in chromosome numbers, speciation events, and extinction events; each of these events occurs at a particular rate (interpreted as the expected number of events per lineage per unit time). The ChromoHiSSE model includes an additional hidden trait with *m* > 1 states. The states of this hidden trait correspond to different sets of ChromoSSE parameters, and lineages evolve among hidden states as a Markov process with an estimated rate.

Under the ChromoHiSSE model, the state of a lineage is both the chromosome number, *n*, and the hidden state, *i*. For numerical stability (Mayrose et al. 2010, Zenil-Ferguson et al. 2017, Freyman & Höhna 2018), we place an upper bound on the possible number of chromosomes (*k*; transitions to *n* > *k* are prohibited, *i*.*e*., have rate 0), and the lower bound is 0. We denote the two hidden states as *i* and *ii*. For example, the state for a lineage could be 10*i*, indicating that the lineage has 10 haploid chromosomes and is in the hidden state *i*.

The ChromoHiSSE model is a stochastic process that begins with two lineages at the root, which evolve independently forward in time. As the process evolves, a lineage can experience anagenetic events (changes in chromosome number or hidden state that happen along the branches of the phylogeny, denoted with subscript _*a*_), cladogenetic events (speciation events that involve changes in the number of chromosomes or hidden state for one of the daughter lineages, hence associated with speciation and denoted with subscript _*c*_), and extinction events. As in all continuous-time Markov chain models, each event happens at an instantaneous point in time and multiple events cannot occur concurrently. The anagenetic events are:

(1) *n* increases by one (*n* + 1 increasing dysploidy) but the hidden state stays the same, which occurs at rate 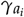;
(2) *n* decreases by one (*n* − 1 decreasing dysploidy) but the hidden state stays the same, which occurs at rate 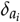, and;
(3) *n* stays the same, but the hidden state changes (*e*.*g*., *i* to *ii*), which occurs at rate *χ*_*a*_. Cladogenetic events produce two daughter lineages that then evolve independently. The states of the daughters depend on the type of event:
(4) both daughters inherit the state of the ancestor, which occurs at rate 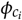;
(5) one daughter inherits *n*, the other inherits *n* + 1 (increasing dysploidy), and both inherit the same hidden state *i*, which occurs with rate 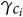;
(6) one daughter inherits *n*, the other inherits *n* − 1 (decreasing dysploidy), and both inherit the same hidden state *i*, which occurs with rate 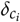;
(7) both daughters inherit *n* from the ancestor, but one daughter changes hidden state (*i* to *ii*), which occurs at rate *χ*_*c*_.

These events are depicted in Fig. 1. Additionally, all lineages go extinct at rate *µ*, independent of *n* or the hidden state. The lineages evolve forward in time until the present, at which point they are sampled independently with probability *f*. Extinct and unsampled lineages are pruned from the tree and the hidden state for sampled lineages is ignored; this produces a realization comprising a reconstructed phylogeny relating the sampled lineages and a chromosome number for each sampled lineage. This stochastic process allows us to compute the probability of an observed dataset (*i*.*e*., the probability that a realization under this process corresponds to our observed data) given a set of parameter values.

**Figure 1.**
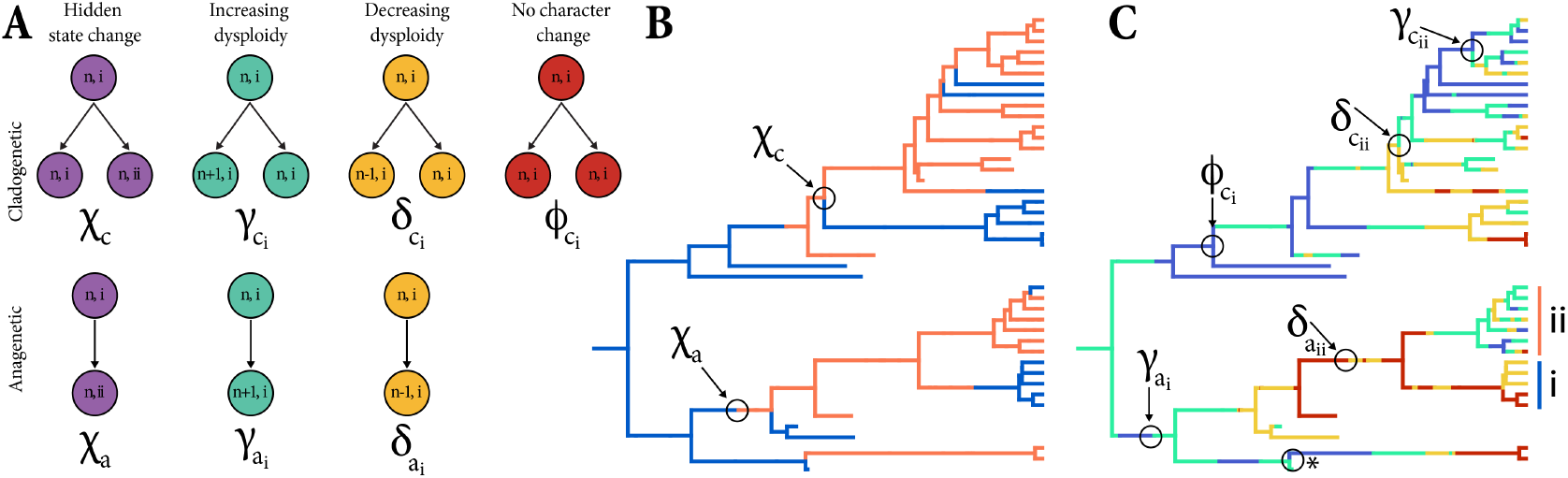
The ChromoHiSSE model. Panel (A) describes the event rates allowed in the model for both cladogenetic (top) and anagenetic (bottom) events. Panels (B) and (C) demonstrate those rates on a tree simulated under ChromoHiSSE. Branches that do not reach the present represent extinct, unsampled lineages. Panel (B) shows the anagenetic and cladogenetic changes in the hidden states, indicated by blue (*i*) vs. orange (*ii*). Panel (C) shows the anagenetic and cladogenetic gains and losses of chromosomes, as well as speciation with no corresponding change in chromosome numbers. More red colors in (C) correspond to more chromosomes. The vertical blue and orange bars in (C) indicate clades in the blue (*i*) vs. orange (*ii*) hidden states, displaying that chromosome number changes are less frequent in the blue hidden state than in the orange. The asterisk in (C) demarcates where a cladogenetic dysploidy event could appear as an anagenetic event because of unsampled lineages.

Note that we assume that there are only two hidden states and that rates of change between hidden states are symmetric; *i*.*e*., the anagenetic rate of change from *i* to *ii* is the same as the rate of change from *ii* to *i*, and likewise for cladogenetic changes. We do not model density-dependent dysploidy rates (rates varying based on the number of chromosomes), as previous studies across all angiosperms that have implemented density-dependent rates have found that the relationship between number of chromosomes and rates of chromosome evolution is very weak to non-existent (Carta et al. 2020). In addition, we assume that the rate of extinction is constant among all lineages, regardless of the number of chromosomes or hidden states. We model a fixed extinction rate because our model is biologically informed by our hypothesis that speciation is linked to chromosome number change, but that dysploid chromosome number changes have little effect on fitness and consequently little effect on extinction. In the supplementary material (S2), we show how to relax some of these assumptions by specifying a full ChromoHiSSE model. These specifications may guide future researchers who wish to include polyploidization (and/or demipolyploidization); similarly, extinction could be allowed to vary in future implementations for empirical scenarios where such variation is warranted.

### Testing differences between transition rates and not states

Most hidden-state SSE (HiSSE) models include two (or more) sets of diversification rates for each state hypothesized to affect diversification (*e*.*g*., Helmstetter et al. 2023). For example, if a researcher wants to know if herbivorous beetles diversify faster than carnivorous ones, the hidden state approach would include two diversification parameters for the carnivorous state and two diversification parameters for the herbivorous state. If both herbivory-specific rates are higher than the carnivory rates, then the analysis suggests that herbivores diversify faster diversification than carnivores. However, our new model differs from previous HiSSE models in that we parameterize diversification rates associated with *types* of state transitions rather than with the states themselves. This means that our goal is not to test whether *n* = 15 or *n* = 16 chromosomes have different modes of diversification, but rather if changes in the karyotype (*i*.*e*., increase: *n* = 15 to *n* = 16 or decrease: *n* = 15 to *n* = 14) are associated with faster or slower diversification rates compared to heterogeneity (or noise) in the diversification process. To do this, we compare speciation rates *not* associated with dysploidy (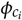 and *χ*_*c*_) to speciation rates *associated* with dysploidy (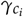 and 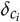) by defining the test statistic 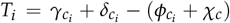 (Fig. 2D). A posterior probability of *T*_*i*_ > 0 greater than 0.95 (*P*(*T*_*i*_ > 0) ≥ 0.95) represents high probability of speciation by single chromosome number change. In contrast, *P*(*T*_*i*_ < 0) ≥ 0.95 suggests that speciation rates unrelated to chromosome number are significantly faster than speciation by single chromosome number change. Test statistic *T*_*ii*_ is defined analogously for the second hidden state.

**Figure 2.**
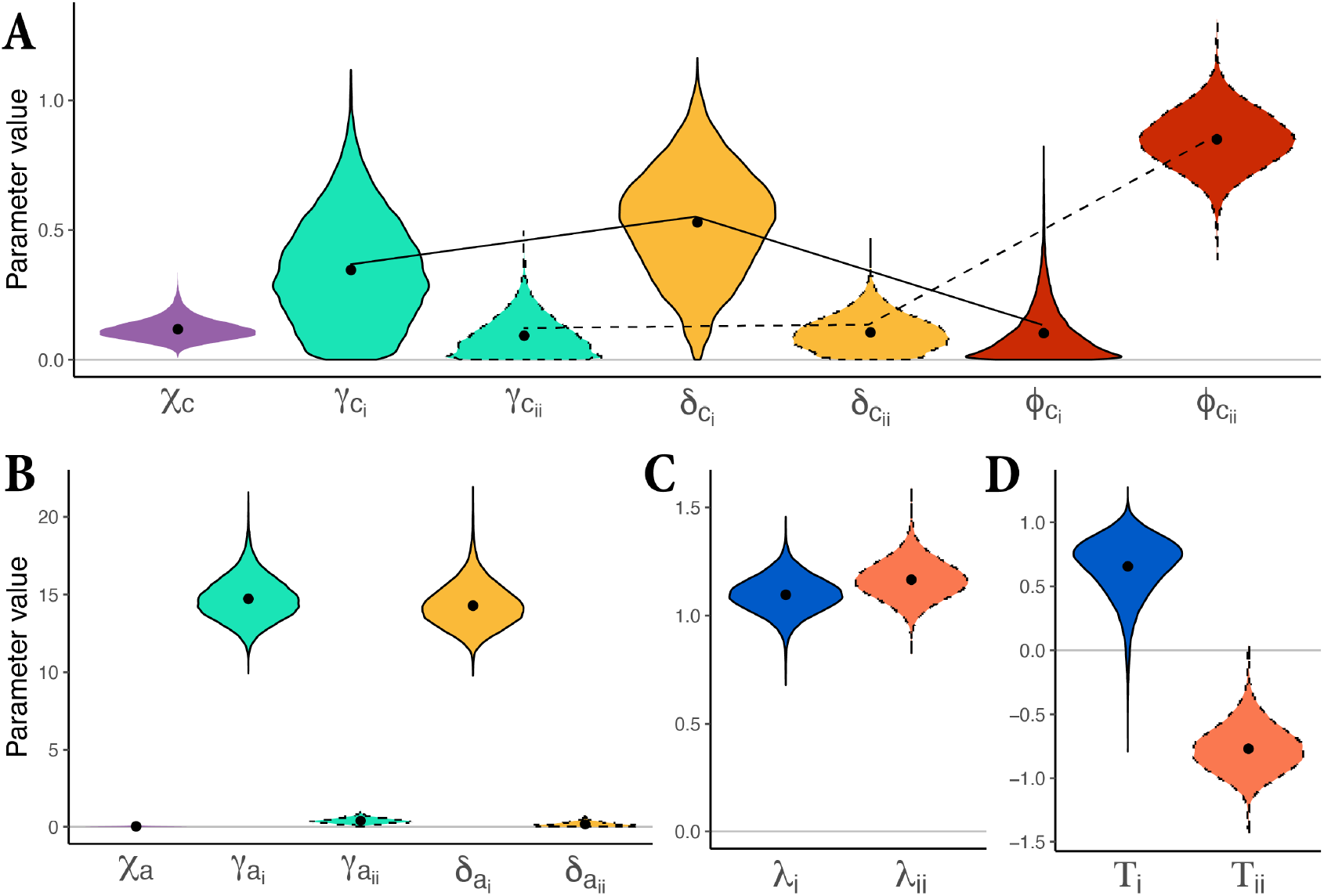
Posterior distributions of rate estimates. Solid lines indicate rates corresponding to hidden state *i* and dashed lines indicate hidden state *ii*. Panel (A) shows the posterior distributions of cladogenetic rates, panel (B) shows the anagenetic rates, panel (C) shows the total speciation rate per hidden state (*λ*), and panel (D) shows the test statistic (*T*_*i*_ and *T*_*ii*_) of the difference in speciation rate associated with chromosome number change and speciation rate without chromosome number change.*T*_*i*_ is significantly greater than 0 (faster speciation when associated with chromosome number change; *P*(*T*_*i*_ > 0) = 0.986) and *T*_*ii*_ is significantly less than 0 (slower speciation when associated with chromosome number change; *P*(*T*_*ii*_< 0) = 1.000)

### Chromosomal and phylogenetic data for *Carex*

We implemented our proposed ChromoHiSSE model on a large dataset that includes a phylogeny of *Carex* with 755 taxa (∼40% of extant diversity), representing all *Carex* subgenera and most of the sections for which chromosome counts have been reported. Our dataset also includes haploid chromosome number (*n*) for all tips in the tree; both the tree and chromosome number data come from (Márquez-Corro et al. 2021). The original tree, based on a HybSeq backbone and three DNA regions (ITS, ETS and *mat*K) from ca. 1400 species out of 2000 *Carex* species, was published by (Martín-Bravo et al. 2019). Tips without chromosome number information were pruned for the current study. Chromosome number in *Carex* evolves through dysploid events, except in the small subgenus *Siderostictae* (13 species in our tree)—sister to the rest of the genus—which includes species with different reported ploidy levels (Márquez-Corro, Martín-Bravo, Spalink, Luceño & Escudero 2019, Márquez-Corro et al. 2021). To avoid modeling rare polyploidy events that occur only in a small part of the tree, we removed *Siderostictae* from the primary analysis (though see Supplemental Section S4 for results with this clade included). For the remaining taxa, we coded chromosome numbers as the most frequent haploid number or the lowest haploid cytotype in polyploid lineages from the Márquez-Corro et al. (2021) dataset for those tips. The final data matrix included *n*-values ranging from 5 to 66 chromosomes for 742 taxa.

### Simulation study

We performed a simulation study to validate ChromoHiSSE using parameter values that approximate our empirical parameter estimates, with the exception of extinction, which we vary across two simulation scenarios. For each scenario (low and high extinction), we simulated 100 datasets (see Supplemental Section S3 for details). We analyzed all datasets using ChromoHiSSE to confirm that our model is able to recover the true simulating values and with ChromoSSE to test for the effect of not accounting for rate variation when it is present in the dataset.

### Computational implementation of ChromoHiSSE

We implemented ChromoHiSSE in RevBayes (Höhna et al. 2016), a software for specifying Bayesian probabilistic graphical models primarily for phylogenetics and phylogenetic comparative methods. Due to the large state space and cladogenesis in our model, we used the newly-developed `TensorPhylo` plugin (May & Meyer 2022) to accelerate likelihood calculations and thus achieve convergence in a more reasonable time frame. We monitored ancestral states through stochastic character mapping as implemented in RevBayes (Freyman & Höhna 2018). We ran two chains of the analysis and assessed convergence in the R programming language (R Core Team 2013). We processed the traces and removed 10% of the generations per chain as burnin using `RevGadgets`. We then calculated the effective sample size (ESS) value for each parameter in each chain using coda (Plummer et al. 2006) and verified that the harmonic mean of the ESS values of each chain was greater than 200. We additionally visually inspected all model parameters across both chains in Tracer (Rambaut et al. 2018). For any parameters that appeared to have strikingly non-normal posterior distributions, we also estimated a transformed ESS following (Vehtari et al. 2021). We subsequently combined both runs for all downstream analyses, discarding 10% of the total generations per chain as burnin prior to combining.

### Analysis and postprocessing

We summarized posteriors and plotted results in R (R Core Team 2013) using the R package `RevGadgets` (Tribble et al. 2022). We additionally transformed model parameters to produce two types of useful summary statistics. First, we calculated the total speciation rate in hidden state *i* vs. *ii* as *λ* = *χ*_*c*_ + *γ*_*c*_ + *δ*_*c*_ + *ϕ*_*c*_. Second, we wanted to know if the rate of speciation concurrent with changes in chromosome number (*γ*_*c*_ + *δ*_*c*_) is greater or less than the rate of speciation with no change in chromosome number (*χ*_*c*_ + *ϕ*_*c*_). We thus estimated an additional summary statistic: *T* = (*γ*_*c*_ + *δ*_*c*_) − (*χ*_*c*_ + *ϕ*_*c*_); positive values of *T* indicate more speciation with chromosome number change and negative values indicate less speciation with chromosome number change. We estimate *T* for each hidden state to estimate how chromosome number changes associated with speciation vary between modes of the model.

All code for implementing and running the model and processing and plotting the results is available at Zenodo DOI: 10.5281/zenodo.8320248.

## Results

### Two modes of chromosomal anagenesis

We recover two modes of anagenetic dysploidy. Branches in hidden state *i* have fast and recurrent single chromosome number increases and decreases (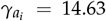 and 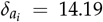 chromosome number events per lineage per million years (E/L/Myr) Table 1, Fig. 2B). For the other branches—in hidden state *ii*—chromosome number rarely changes via dysploidy (increases are 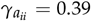 and decreases are 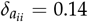 E/L/Myr, Table 1, Fig. 2B). Rarely, lineages transition between hidden states (*χ*_*a*_ = 0.02). In Table 1 we show the credible intervals for these estimates, and in figure 2B we show the posterior distribution of all the anagenetic rate estimates.

**Table 1.** Summary statistics of posterior distributions for parameter estimates for the ChromoHiSSE model. Light blue rows correspond to parameter estimates for hidden state *i*, and light orange rows correspond to parameter estimates for hidden state *ii*. All rate estimates are given in units of events per lineage per million years (E/L/Myr)

|  | Model parameter | Parameter Type | Hidden State | Median | 2.5% Quantile | 97.5% Quantile |
| --- | --- | --- | --- | --- | --- | --- |
| $\chi_c$ | Hidden state change | Cladogenetic | $i, ii$ | 0.113 | 0.051 | 0.215 |
| $\gamma_{c_i}$ | Increasing dysploidy | Cladogenetic | $i$ | 0.330 | 0.021 | 0.772 |
| $\gamma_{c_{ii}}$ | Increasing dysploidy | Cladogenetic | $ii$ | 0.077 | 0.003 | 0.261 |
| $\delta_{c_i}$ | Decreasing dysploidy | Cladogenetic | $i$ | 0.539 | 0.120 | 0.916 |
| $\delta_{c_{ii}}$ | Decreasing dysploidy | Cladogenetic | $ii$ | 0.099 | 0.007 | 0.250 |
| $\phi_{c_i}$ | No character change | Cladogenetic | $i$ | 0.072 | 0.003 | 0.363 |
| $\phi_{c_{ii}}$ | No character change | Cladogenetic | $ii$ | 0.849 | 0.642 | 1.058 |
| $\chi_a$ | Hidden state change | Anagenetic | $i, ii$ | 0.021 | 0.001 | 0.093 |
| $\gamma_{a_i}$ | Increasing dysploidy | Anagenetic | $i$ | 14.638 | 12.282 | 17.626 |
| $\gamma_{a_{ii}}$ | Increasing dysploidy | Anagenetic | $ii$ | 0.397 | 0.121 | 0.769 |
| $\delta_{a_i}$ | Decreasing dysploidy | Anagenetic | $i$ | 14.191 | 11.903 | 17.246 |
| $\delta_{a_{ii}}$ | Decreasing dysploidy | Anagenetic | $ii$ | 0.142 | 0.005 | 0.505 |

### Two modes of chromosomal evolution and speciation

We identify variation in modes of chromosomal evolution across the tree via two distinct modes, corresponding to the two hidden states *i, ii*. While the overall total speciation rate (*λ*) between the two modes is very similar (Fig. 2C), rates of dysploidy and dysploidy-driven speciation vary significantly between the modes.

Hidden state *i* is characterized by a positive association between speciation and dysploidy and generally high rates of cladogenetic dysploidy (Fig. 2, solid lines, and Table 1). In this mode, the difference between the rate of speciation with chromosome number change and speciation without chromosome number change is larger than zero with 98.6% probability (*P*[*T*_*ii*_>0.986). Thus in hidden state *i*, there is a highly probable and positive correlation between chromosome number change and speciation rates (Fig. 2D, blue). Every estimated dysploidy-related cladogenetic rate is higher than non-dysploidy rates (*e*.*g*., speciation with no state change, Fig. 2A), which suggests that speciation happens more frequently when associated with changes to karyotype.

Cladogenetic increasing and decreasing dysploidy (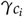 and 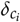) are higher than 0 with 99% probability (Table 1) and faster than the cladogenetic rate of no character change (speciation without change in chromosome number, 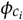). While 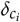 is slightly faster than 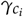, these rates overlap (Table 1).

In contrast, hidden state *ii* is characterized by a negative association between speciation and dysploidy and slower rates of anagenetic and cladogenetic dysploidy (Fig. 2, dashed lines, and Table 1). The difference between speciation associated with chromosome number change and speciation without chromosome number change is less than 0 E/L/Myr (*P*(*T*_*ii*_ < 0) = 1.000), suggesting that speciation is slower when associated with a chromosome number change in hidden state *ii* (Fig. 2D, orange). Anagenetic rates (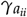 and 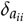) are quite low, approximating 0 (Table 1, Fig. 2B), suggesting that little to no anagenetic change in chromosome number happens in hidden state *ii*.

Cladogenetic dysploidy rates are also quite low, and while the rate of decreasing dysploidy 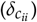 is slightly higher than increasing dysploidy 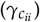, this difference is minimal (Table 1). In summary, we identify variation in modes of chromosomal evolution across the tree and via two distinct modes, corresponding to the two hidden states *i, ii*. In hidden state *i* dysploidy is linked to speciation, whereas in hidden state *ii*, with the same total speciation rate, dysploidy is not an important contributor to this speciation rate.

### Reconstruction of chromosome number evolution

Just over half (51.85%) of all branches in the phylogeny showed a net change in chromosome number (Fig. 3). Overall, the dominant pattern of chromosome number in *Carex* is one of frequent but gradual change; 75.05% of per-branch net chromosome number changes are three or fewer gains or losses. Most clades vary in chromosome number despite some shallow evolutionary divergences (Fig. 3A). Yet there are some remarkable exceptions. For example, clade 2— which includes sect. *Pictae*, the Hirta Clade, and sect. *Praelongae*— shows substantial variation in chromosome number, from *n* = 13 to *n* = 66. However, a clade of hooked sedges (*Carex* sect. *Uncinia*; Fig. 3, clade 1 in grey), has a constant chromosome number (*n* = 44) across the 33 species included in this study, with one exception: *Carex perplexa* has a count of *n* = 66, suggesting that this species may be a demipolyploid. While both clades highlighted in Fig. 3 have similar crown ages, number of species, and average chromosome number (*n* = 35.7 vs. *n* = 44.6), they differ dramatically in karyotype variability and, consequently, in the inferred hidden state (clade 1 = *ii*, clade 2 = *i*, Fig. 3C).

**Figure 3.**
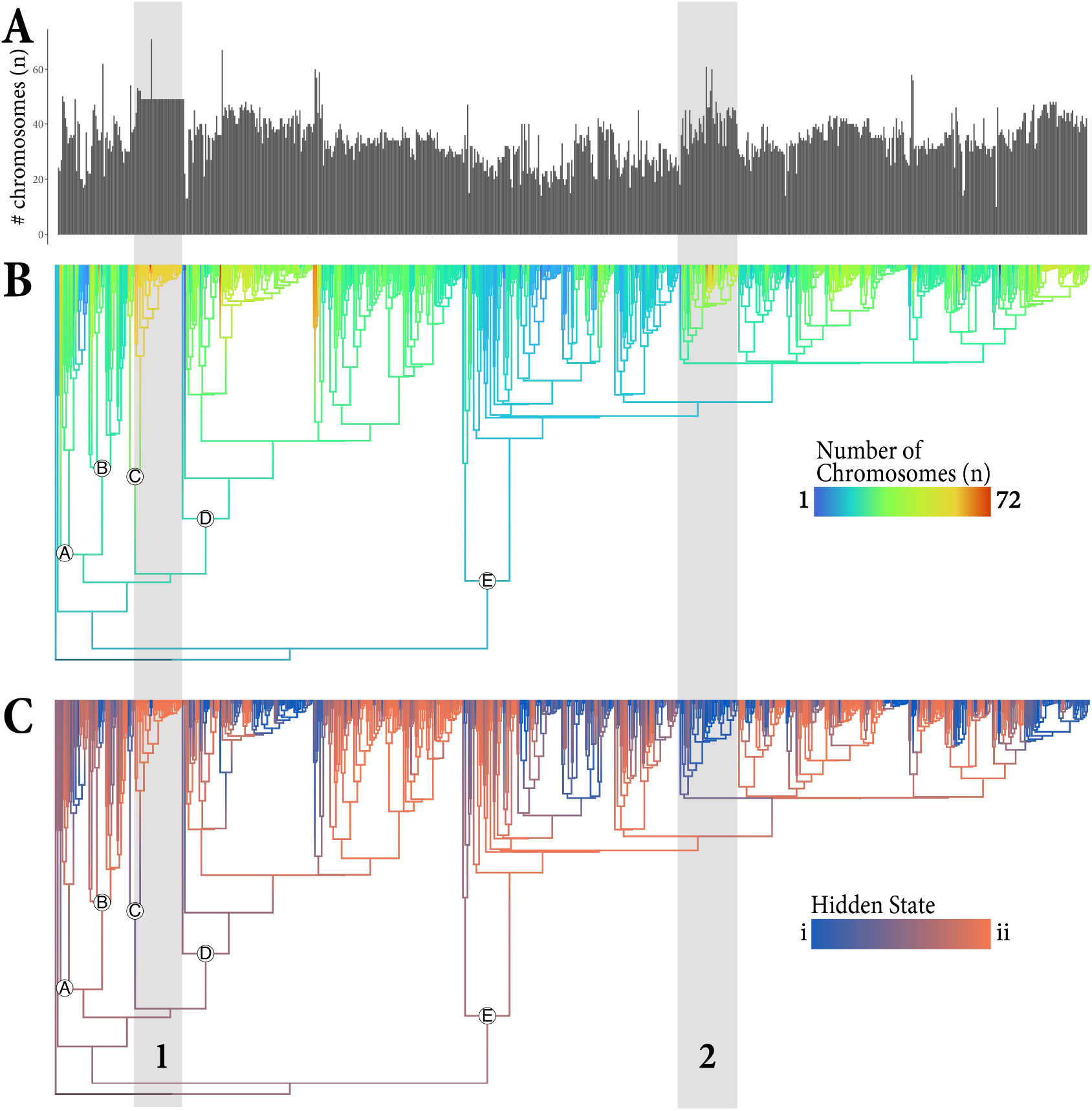
Reconstruction of chromosome numbers (A and B) and hidden states (C) on the *Carex* phylogeny. Panel (A) shows the distribution of haploid chromosome numbers for all extant taxa included in the analysis. Panel (B) shows the reconstructed evolution of chromosome number along branches of the phylogeny, where lighter colors indicate more chromosomes. Panel (C) shows the reconstructed evolution of the hidden state along branches of the phylogeny, where blue indicates strong statistical support for state *i*, orange indicates strong support for state *ii*, and intermediary colors indicate uncertainty in the estimates. Grey bars highlight two clades (labeled 1 and 2) that are discussed in the main text, and circled letters on nodes indicate the position of major subgenera: A = *Psyllophora*, B = *Euthyceras*, C = *Uncinia*, D = *Vignea*, E = *Carex*.

### Simulation study

Our ChromoHiSSE analyses of datasets simulated under ChromoHiSSE (with low and high extinction scenarios) demonstrate that our model generally recovers the true, simulating parameter values, with three exceptions, discussed below. In contrast, when we analyze data simulated under ChromoHiSSE with a ChromoSSE model, parameter estimates are severely compromised.

For analyses under ChromoHiSSE, our simulations with a high extinction rate reveal a tendency to underestimate anagenetic dysploidy rates for the hidden state with the highest rates of dysploidy (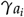 and 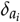), as characterized by the error in the posterior mean point estimate (squared error of the posterior mean normalized by the true value averaged across simulations of ≈0.6 and ≈0.55 for the two parameters, respectively) as well as the frequency with which the 95% credible interval of the posterior contains the true value (the “coverage”; ≈26% and ≈32% for the two parameters, respectively). However, the difference between these rates 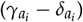 is less biased (normalized squared error of ≈28) and the coverage is nominal (≈95%). The tendency to underestimate high rates could be a result of saturation (so many chromosome number changes that subsequent changes become “invisible” in the simulated datasets) or prior sensitivity (as our prior mean on these rates is significantly lower than the true value, concentrating the prior probability on low rates; see prior specification details in Supplemental Section S3). Additionally, the posterior mean rates of change between hidden states (both anagenetic *χ*_*a*_ and cladogenetic *χ*_*c*_) are higher than the simulating values (Figs. S1 and S2). However, the coverage for these parameters is high (> 95%), suggesting that our ability to detect the true rate of change between hidden states is low. All of these results are similar for high-extinction and low-extinction simulation scenarios, and are presented more detail in Supplemental Section S3.

For analyses under ChromoSSE, the posterior distributions are intermediate between the true values for the hidden states, suggesting that ChromoSSE is essentially “averaging” the rates for the two hidden states together. Additionally, the “95%” credible intervals of these distributions do not contain the true value of either hidden state. (See Supplementary Material, Figures S3 and S6, Tables S3 and S5 for detailed results under the ChromoSSE model.)

## Discussion

Although the impact of dysploidy on *Carex* diversification has been addressed previously (*e*.*g*., Faulkner 1972, Márquez-Corro, Martín-Bravo, Pedrosa-Harand, Hipp, Luceño & Escudero 2019, Márquez-Corro et al. 2021, Hipp 2007), this is the first study to jointly model chromosome number change and diversification, and to demonstrate an association between higher speciation rates and dysploidy in parts of the phylogeny despite heterogeneity in the process of diversification. While gains and losses in chromosome number spur diversification along parts of the phylogeny (hidden state *i*), they have the opposite effect elsewhere in the tree (hidden state *ii*, see Fig. 3). Furthermore, while dysploidy does not lead to higher rates of speciation across the tree on average, speciation in some clades is driven strongly by dysploidy. We propose that these discrepancies between clades may be due to the nature of holocentric chromosomes, where a single dysploidy event in isolation may not be enough to trigger reproductive isolation (Hipp et al. 2010, Lucek et al. 2022, Márquez-Corro, Martín-Bravo, Pedrosa-Harand, Hipp, Luceño & Escudero 2019, Escudero, Hahn, Brown, Lueders & Hipp 2016, Whitkus 1988), but the accumulation of sufficient chromosomal rearrangements in a lineage may form a reproductive barrier and thus trigger speciation (Escudero, Hahn, Brown, Lueders & Hipp 2016, Whitkus 1988, Baker & Bickham 1986). This is a central hypothesis discussed in Fig. 4 of Lucek et al. (Lucek et al. 2022) known as the recombination suppression/hybrid dysfunction chromosomal speciation model. Below, we elaborate on the differences between speciation modes, discuss the evidence supporting the recombination suppression/hybrid dysfunction model, and provide suggestions for future research.

Some *Carex* species are known to have striking chromosome-number polymorphism, even within populations (Luceño & Castroviejo 1991, Escudero, Arroyo, González-Ramírez & Jordano 2023, Escudero et al. 2013, Whitkus 1988). For example, *Carex scoparia* varies from *n* = 28 to *n* = 35 (Escudero et al. 2013), and individuals with different chromosome numbers are able to reproduce and exchange alleles, maintaining gene flow despite chromosome number differences. This polymorphism suggests that some chromosome number differences are insufficient to create reproductive barriers (Hipp et al. 2009), which is supported by our results; often, inferred chromosome number changes do not result in an immediate speciation event on the phylogeny. This constant gene flow might also result in a difficulty to estimate anagenetic chromosome number change when is rampant. As our simulations have indicated, it is possible that anagenetic dysploidy is underestimated frequently for holocentric chromosomes. The question remains, why does dysploidy result in speciation in some cases and not in others? One possibility is that an accumulation of changes may eventually lead to reproductive isolation, where one “last straw” dysploidy event triggers speciation (referred to as the last-straw hypothesis). We discuss potential model developments that could test this theory below in the *Modeling chromosomal speciation* section. Another possibility is that rearrangements in some parts of the genome are more stable than others, and the genomic architecture—where the fragmentation or fusion occurs in the genome—determines the evolutionary effects of dysploidy.

Patterns of repeat DNA—*e*.*g*., LINEs, LTRs, and Helitrons—differ significantly between *Carex* lineages whose chromosome arrangements evolve at different rates (Cornet et al. 2023). Moreover, repeat regions in *Carex* genomes investigated correlate with chromosome breakpoints across deep phylogenetic breaks as well as within species (Escudero, Marques, Lucek & Hipp 2023, Höök et al. 2023), pointing to a mechanism by which natural selection could operate on the genome architecture of chromosome rearrangements in the genus. In fact, synteny in holocentric sedge chromosomes is more conserved than we would expect if chromosome evolution were unconstrained, even in comparisons between species that span deep nodes in the phylogeny (Escudero, Marques, Lucek & Hipp 2023). This suggests that selection may maintain large blocks of the genome, likely comprising numerous, tightly linked genes (Escudero, Marques, Lucek & Hipp 2023). The massive diversity of sedges is thus likely an outcome of two processes: recombination suppression in rearranged regions of the genome, shaping ecological divergence in during species divergence; and the typically-gradual—or, occasionally, immediate—evolution of reproductive isolation as chromosomes split and fuse. Such information could be incorporated into a macroevolutionary framework by relaxing the assumption in SSE models that speciation occurs instantly, and instead integrating microevolutionary dynamics of how genomic changes become fixed in lineages via ancestral recombination graphs (*e*.*g*., Deng et al. 2024, Brandt et al. 2024).

An important caveat to our study is the presence of missing data—particularly for tropical lineages—both in terms of sequenced taxa represented in the phylogeny and for available chromosome counts. While the phylogeny used in this study was assembled with a HybSeq backbone and three DNA regions for ca. 1400 out of 2000 species, phylogenetic relationships in some areas of the genus are still tenuous (Martín-Bravo et al. 2019, Roalson et al. 2021, Jiménez-Mejías et al. 2016). Additionally, there are only chromosome counts for ∼ 700 species in the phylogeny, and data availability is more sparse in tropical lineages than temperate ones. For example, one of the most species-diverse lineages within *Carex*—the Decora Clade/*Carex* sect. *Indicae* (Roalson et al. 2021)—only includes five species with reported chromosome numbers (Márquez-Corro et al. 2021). This pantropical lineage alone would constitute ca. 10% of the genus (Roalson et al. 2021). Also, we include only a few New Zealand species from *Carex* sect. *Uncinia* (clade 1 in Fig. 3), as there are few known karyotypes for the South American representatives of the lineage. Dispersal traits such as hooked utricles and epizoochory might be as or more important for diversification as karyotype in this clade, but we cannot tease these effects apart without additional chromosome count data. *Carex* biodiversity is greatest in temperate regions including the Global North, and this bias in data availability may suggest that our findings are most robust for northern temperate *Carex*. Filling these data gaps would facilitate testing whether evolutionary modes of dysploidy play out differently based on geography and/or dispersal traits.

### Chromosome number evolution

The contrasting clades 1 and 2 from Fig. 3 reflect the two modes of macroevolution described above in “Results: Reconstruction of chromosome number evolution” and in “Discussion: Multiple modes of chromosomal speciation” exemplify the importance of including hidden states in our model. *Carex* sect. *Uncinia* (hook sedges, clade 1 in Fig. 3) is characterized by hooked utricles that allow for long-distance dispersal through epizoochory (García-Moro et al. 2022). The hook sedges exemplify a *Carex* clade that is characterized by low-to-zero dysploidy but high rates of speciation (Martín-Bravo et al. 2019)—though this was not formally tested in our study—and thus may be a particularly appealing candidate for future studies focusing on potential adaptive traits (currently “hidden” and not related to chromosome number) that drive diversification. This clade is characterized by similar karyotypes of many (*n* = 44) chromosomes (Márquez-Corro et al. 2021), mostly reported from New Zealand, where ca. half of the section diversified. However, this only gives partial information on their evolutionary history, as the karyotypes of the Andean relatives have not been studied in detail yet. Most of the New Zealand chromosome counts correspond to a single, old report and there is some variation in the few existing counts from South America, so these results may be taken with caution. If the South American species show wider karyotypic variation than currently detected in comparison to the New Zealand lineage, there may be different drivers enhancing speciation on each side of the South Pacific.

### Modeling chromosomal speciation

The methodological innovations presented here allow us to integrate process noise and variation in chromosome number evolution, despite the computational challenges associated with the huge number of states defined by chromosome numbers. Process variation is fundamental to our conclusions; only by incorporating multiple diversification modes do we discover that while dysploidy is strongly associated with speciation in some clades, in other parts of the phylogeny, dysploidy is rare and does not lead to faster speciation. Leaving the hidden states out leads to a false inference about the significance of chromosome evolution to diversification: when we implement our model without process variation, our results suggest a strong, uniform boost in speciation rate associated with dysploidy (results presented in Supplemental Section S4). Furthermore, our simulation study demonstrates that when hidden rate variation is present, analyses by a model without hidden rate variation (*e*.*g*., ChromoSSE) consistently fail to recover the true parameter values (Supplemental Fig. S3 and Fig. S6). Our approach permits reconstructions of the past via stochastic maps, allowing for the detection of lineages in which dysploidy has been linked or not to diversification. Reconstruction of the past and identification of clade and lineages where a trait makes a difference in diversification is key for future smaller comparative studies aiming to understand the genomic underpinnings of plant speciation.

RevBayes’ graphical modeling framework permits flexibility in future modifications to our approach. Our ChromoHiSSE model does not include parameters for polyploidy, but we provide the mathematical framework for such additions in Supplemental Section S2. Our model includes two hidden states, but future implementations could increase the number of hidden states (with a significant increase in the computational effort required) and/or limit which parameters vary across hidden states. Model selection—typically quite challenging and computationally intensive to implement in Bayesian approaches—could assist researchers with smaller datasets, who lack the ability to estimate all parameters in this parameter-rich model, to decide between models that allow all or some parameters to vary between hidden states. In particular, we believe a Bayesian model averaging approach using reversible-jump MCMC may prove particularly useful (Freyman & Höhna 2018), as it avoids the need to compute computationally expensive marginal likelihoods, and could automatically consider all ways that parameters could be shared between hidden states.

Future work that builds off of our ChromoHiSSE approach will allow us to pursue promising avenues for innovative research; we highlight two examples below. First, like all birth-death models, our ChromoHiSSE model operates with species as the fundamental unit of analysis (the tips in the tree) and thus does not formally model chromosome number polymorphism that is present in some *Carex* lineages. Second, while ChromoHiSSE tests for the effect of single changes in chromosome number on diversification rates, it cannot test for the effect of an accumulation of changes (the last-straw hypothesis). However, ChromoHiSSE could be modified to include tip-state polymorphism (*e*.*g*., a dysploid series) as additional hidden states (*e*.*g*., a particular tip either has 12, 13, or 14 chromosomes). Additionally, (Goldberg & Foo 2020) described a mechanism for modeling memory (thus the accumulation of chromosome number changes) in a macroevolutionary framework using hidden states, which could be applied to test the last-straw hypothesis.

## Conclusions

Our work demonstrates the important role of dysploidy on diversification in *Carex*, a model lineage for understanding how karyotype rearrangements via dysploidy affect speciation and macroevolutionary dynamics. Our results—using the new ChromoHiSSE model—paint a complex picture of how dysploidy affects speciation in a clade characterized by high species diversity, high morphological disparity, and holocentric chromosomes (Martín-Bravo et al. 2019), and our results support the recombination suppression/hybrid dysfunction chromosomal speciation model, in which only some karyotype rearrangements trigger reproductive isolation and thus speciation. Future work on the underlying genomic mechanisms of chromosomal speciation via comparative genomics will be particularly powerful for linking across scales, from molecules to lineages. Ultimately, our novel modeling approach also serves as a critical step towards even more complex and powerful macroevolutionary analyses that incorporate intraspecific chromosome number variation and track the accumulation of change through time.

## Author Contributions

RZF and ME conceived the study. CMT, JIM-C, ME, and RZF wrote the article. JIM-C, ME, ALH gathered all *Carex* chromosome number and phylogenetic data. CMT, MRM, and RZF designed models and methods, and verified mathematical/ statistical notation. CMT and MRM wrote computational code and created figures. All authors revised and edited the manuscript.

## Data availability

RevBayes model code, and R code for processing and plotting the results are available at Zenodo DOI: 10.5281/zenodo.8320249

## Acknowledgements

CMT was supported by the School of Life Sciences at the University of Hawai’i at Mānoa funded through the Office of the Vice President of Research. This material is based upon work supported by the NSF Postdoctoral Research Fellowships in Biology Program under Grant No. 2109835 to CMT. Any opinions, findings, and conclusions or recommendations expressed in this material are those of the authors and do not necessarily reflect the views of the National Science Foundation. JIM-C was granted by the ‘Next Generation EU’ funding, the Recovery Plan, Transformation and Resilience and the Ministry of Universities, under the grants ‘Margarita Salas’ for the requalification of the Spanish university system 2021–2023 called by the Universidad Pablo de Olavide, Seville. ME was supported by the MICINN-FEDER, Project DiversiChrom (PID2021-122715NB-I00). RZF was supported by NSF-DEB 2323170. We thank three anonymous reviewers for their comments and feedback on earlier versions of the manuscript.

## Supporting Information

**Supporting Information 1**. Detailed description of the Chromosome number and Hidden State-dependent Speciation and Extinction model (ChromoHiSSE).

