## Supplemental Material for "Macroevolutionary inference of complex modes of chromosomal speciation in a cosmopolitan plant lineage"

#### S1 Background on dysploidy and diversification

Most studies of chromosomal rearrangements (dysploidy or inversions) focus on population-level samples (within a single species) or interspecific hybrids (*e.g.*, Carr, 1975; Livingstone et al., 2000; Rieseberg, 2001; Navarro and Barton, 2003). Therefore, the macroevolutionary scale-level data necessary for testing the role of dysploidy across phylogenies is often lacking (Escudero et al., 2014; Mandáková and Lysak, 2018). Additionally, theoretical studies disagree over the importance of dysploidy in diversification. For example, in an interesting modeling and simulation study, Guerrero and Kirkpatrick (2014) provided evidence that chromosomal fusions (decreasing dysploidy) can have adaptive advantages due to reduced recombination of locally adapted alleles. Alternatively, dysploidy could contribute to reproductive isolation in populations and lead to sympatric speciation through either hybrid dysfunction or recombination suppression (*e.g.*, Rieseberg, 2001). In the hybrid-dysfunction model, there is a strong selection against heterozygotes (underdominance) from crosses between parents with different karyotypes (Coyne and Orr, 2004). Hipp et al. (2009) suggested that hybrid dysfunction may explain speciation patterns in *Carex*. The model suffers a theoretical challenge: the probability of fixation of new chromosomal variants is very small, as the individuals that carry a new karyotype have a low chance of interbreeding with other individuals in the population and becoming established. New karyotypes may be maintained only if selection against karyotypic heterozygotes is weak. But in such cases, reproductive isolation would also be minimal, reducing the probability of rearrangement leading to speciation. This is known as the underdominance paradox (Spirito, 1998). Because of this paradox, the hybrid-dysfunction model has been criticized (Ayala and Coluzzi, 2005). An alternative, the recombination-suppression model of chromosomal speciation, has gained more acceptance in the last two decades (Butlin, 2005). In this model, locally adapted genes are clustered in supergenes where there is no recombination because of structural incompatibilities, typically an inversion or a fission-fusion rearrangement. Chromosomal speciation may consequently happen despite gene flow between the ancestral and the novel karyotype. Lucek et al. (2022) discussed the hybrid dysfunction and recombination suppression models in the context of species with holocentric chromosomes (defined below), arguing that these classic speciation models were built around the idea that chromosomal rearrangements fixed between species cause hybrid sterility in monocentrics, due to unbalanced meiotic products or segregation problems in heterozygotes.

Chromosome fissions, fusions, and inversions have great evolutionary potential. Chromosome fusions not only reduces recombination between formerly unlinked genes but may also reduce recombination rates within each arm (Dumas and Britton-Davidian, 2002). Fusions could be favored in situations when the tight linkage is beneficial (Charlesworth, 1985). Recombination is suppressed in heterozygous karyotypes that have experienced an inversion event, and so the locally adapted alleles are bound together in a single high-fitness chromosome. By reducing recombination, inversions preserve the divergence between populations in the presence of gene flow, setting the stage for potential speciation (Navarro and Barton, 2003; Kirkpatrick and Barton, 2006).

#### S2 The General ChromoHiSSE Model

Here we present the general ChromoHiSSE model, which includes additional chromosome-change events from ChromoSSE (Freyman and Höhna, 2018), an arbitrary number of hidden states, and arbitrary rates of change between pairs of hidden states. As a reminder, in the ChromoHiSSE model we fitted for *Carex* in the main manuscript the anagenetic and cladogenetic rates we estimated were:

(1) $n$ increases by one ($n+1$ increasing dysploidy) but the hidden state stays the same, which occurs at rate $\gamma_{a_i}$;

(2) $n$ decreases by one ($n-1$ decreasing dysploidy) but the hidden state stays the same, which occurs at rate $\delta_{a_i}$, and;

(3) $n$ stays the same, but the hidden state changes ($i$ to $ii$), which occurs at rate $\chi_a$.

(4) both daughters inherit the state of the ancestor, which occurs at rate $\phi_{c_i}$;

(5) one daughter inherits $n$, the other inherits $n+1$ (increasing dysploidy), and both inherit the same hidden state $i$, which occurs with rate $\gamma_{c_i}$;

(6) one daughter inherits $n$, the other inherits $n-1$ (decreasing dysploidy), and both inherit the same hidden state $i$, which occurs with rate $\delta_{c_i}$;

(7) both daughters inherit $n$ from the ancestor, but one daughter changes hidden state ($i$ to $ii$), which occurs at rate $\chi_c$.

The additional anagenetic events in the general ChromoHiSSE model may include (continuing from the list in the main text):

(8) $n$ increases to $2n$ (polyploidization), which occurs at rate $\rho_{a_i}$;

(9) for even $n$, $n$ increases to $1.5n$ (demi-polyploidization), which occurs at rate $\eta_{a_i}$;

(10) for odd $n > 2$, $n$ increases to $\lceil 1.5n \rceil$ at rate $0.5\eta_{a_i}$ and increases to $\lfloor 1.5n \rfloor$ at rate $0.5\eta_{a_i}$.

and in addition it may include the following cladogenetic changes:

(10) one daughter inherits $n$, the other inherits $2n$ (polyploidization), and both inherit the hidden state, which occurs with rate $\rho_{c_i}$;

(11) for even $n$, one daughter inherits $n$, the other inherits $1.5n$ (demi-polyploidization), and both inherit the hidden state, which occurs with rate $\eta_{c_i}$;

(12) for odd $n > 2$, one daughter inherits $n$, the other inherits $\lceil 1.5n \rceil$ (demi-polyploidization), and both inherit the hidden state, which occurs with rate $0.5\eta_{c_i}$;

(13) for odd $n > 2$, one daughter inherits $n$, the other inherits $\lfloor 1.5n \rfloor$ (demi-polyploidization), and both inherit the hidden state, which occurs with rate $0.5\eta_{c_i}$.

Furthermore, we allow there to be $m$ hidden states labeled arbitrarily.

In the most general ChromoHiSSE model, rates of change between pairs of hidden states are allowed to vary. The two parameters that require modification to allow hidden state rates to transition asymmetrically are: , such that the state-change events are (modifying list elements in the main text):

(3) (anagenetically) $n$ stays the same, but the hidden trait changes from state $i$ to state $ii$, which occurs at rate $\chi_{a_{i,ii}}$;

(7) (cladogenetically) both daughters inherit $n$ from the ancestor, but one daughter changes from the ancestral hidden state $i$ to hidden state $ii$, which occurs at rate $\chi_{c_{i,i,ii}}$ (where the first subscript refers to the ancestor, and the second and third refer to the daughters).

Additionally, we define the total anagenetic and cladogenetic rates of leaving hidden state $i$ as $\chi_{a_i} = \sum_{ii \neq i}^{m} \chi_{a_{i,ii}}$ and $\chi_{c_i} = \sum_{ii \neq i}^{m} \chi_{c_{i,i,ii}}$, respectively.

Finally, we allow the rate of extinction to depend on the hidden state, denoted $\mu_i$.

The model presented in the main text is a special case of the general ChromoSSE, we are simplifying it by assuming that:

(1) Polyploidy and demiploidy rates are zero both at anagenesis and cladogenis. Therefore, $\rho_{a_i} = \eta_{a_i} = \rho_{c_i} = \eta_{c_i} = 0$.

(2) Anagenetic hidden rates are symmetric $\chi_{a_{i,ii}} = \chi_{a_{ii,i}}$ for any two hidden states $i, ii$, and cladogenetic hidden rates are symmetric as well $\chi_{c_{i,i,ii}} = \chi_{c_{ii,ii,i}}$ for all $i, ii$ for any two hidden states.

(3) Extinction rates for all hidden states are the same $\mu_i = \mu_{ii}$ for all $\{i, ii\}$.

(4) The number of hidden states is two, that is $m = 2$.

We compute the probability of the reconstructed tree and chromosome counts (*i.e.*, the likelihood) using standard calculations for SSE models (Maddison et al., 2007). As in Maddison et al. (2007), we can write the stochastic differential equations(SDEs, Kolmogorov forward) to describe how extinction probabilities—denoted $E_{n,i}(t)$—and lineage probabilities—denoted $D_{l,n,i}(t)$ for lineage $l$—change over time; we present these SDEs below.

#### S2.1 Extinction probabilities

The quantity $E_{n,i}(t)$ represents the probability that a lineage alive at time $t$ with $n$ chromosomes and in hidden state $i$ leaves no sampled descendants at the present (either because of extinction or incomplete sampling). This quantity is necessary to accommodate missing speciation events along branches of the phylogeny. We derive the stochastic differential equations that describe how the extinction probability changes over time following standard techniques for SSE models (Maddison et al., 2007; Goldberg and Igić, 2012).

The opportunity for different kinds of cladogenetic events depends on the number of chromosomes; *e.g.*, decreasing disploidy is impossible when $n = 1$.

The SDEs for all cases are:

$$
\begin{aligned}
\frac{dE_{n,i}(t)}{dt} = \mu_i - &\left[\gamma_{a_i} + \rho_{a_i} + \eta_{a_i} + \chi_{a_i} + \phi_{c_i} + \gamma_{c_i} + \rho_{c_i} + \eta_{c_i} + \chi_{c_i} + \mu_i\right] E_{n,i}(t) + \\
&\left[\gamma_{a_i} + \rho_{a_i}\right] E_{n+1,i}(t) + \sum_{ii \neq i}^{m} \chi_{a_{i,ii}} E_{n,ii}(t) + \\
&\phi_{c_i} E_{n,i}(t)^2 + \left[\gamma_{c_i} + \rho_{c_i}\right] E_{n+1,i}(t) E_{n,i}(t) + \sum_{ii \neq i}^{m} \chi_{c_{i,i,ii}} E_{n,i}(t) E_{n,ii}(t) \qquad \text{for } n = 1
\end{aligned}
\tag{S1}
$$

$$
\begin{aligned}
\frac{dE_{n,i}(t)}{dt} = \mu_i - & \left[\gamma_{a_i} + \delta_{a_i} + \rho_{a_i} + \eta_{a_i} + \chi_{a_i} + \phi_{c_i} + \gamma_{c_i} + \delta_{c_i} + \rho_{c_i} + \eta_{c_i} + \chi_{c_i} + \mu_i\right] E_{n,i}(t) + \\
& \gamma_{a_i} E_{n+1,i}(t) + \delta_{a_i} E_{n-1,i}(t) + \rho_{a_i} E_{2n,i}(t) + \eta_{a_i} E_{1.5n,i}(t) + \sum_{ii \neq i}^{m} \chi_{a_{i,ii}} E_{n,ii}(t) + \\
& \phi_{c_i} E_{n,i}(t)^2 + \gamma_{c_i} E_{n+1,i}(t) E_{n,i}(t) + \delta_{c_i} E_{n-1,i}(t) E_{n,i}(t) + \eta_{c_i} E_{1.5n,i}(t) E_{n,i}(t) + \\
& \rho_{c_i} E_{2n,i}(t) E_{n,i}(t) + \sum_{ii \neq i}^{m} \chi_{c_{i,i,ii}} E_{n,i}(t) E_{n,ii}(t) \qquad \text{for } 2 < n < k/2, n \text{ even}
\end{aligned}
\tag{S2}
$$

$$
\begin{aligned}
\frac{dE_{n,i}(t)}{dt} = \mu_i - & \left[\gamma_{a_i} + \delta_{a_i} + \rho_{a_i} + \eta_{a_i} + \chi_{a_i} + \phi_{c_i} + \gamma_{c_i} + \delta_{c_i} + \rho_{c_i} + \eta_{c_i} + \chi_{c_i} + \mu_i\right] E_{n,i}(t) + \\
& \gamma_{a_i} E_{n+1,i}(t) + \delta_{a_i} E_{n-1,i}(t) + \rho_{a_i} E_{2n,i}(t) + 0.5\eta_{a_i} \left[E_{\lceil 1.5n \rceil,i}(t) + E_{\lfloor 1.5n \rfloor,i}(t)\right] + \\
& \sum_{ii \neq i}^{m} \chi_{a_{i,ii}} E_{n,ii}(t) + \phi_{c_i} E_{n,i}(t)^2 + \gamma_{c_i} E_{n+1,i}(t) E_{n,i}(t) + \delta_{c_i} E_{n-1,i}(t) E_{n,i}(t) + \\
& \rho_{c_i} E_{2n,i}(t) E_{n,i}(t) + 0.5\eta_{c_i} \left[E_{\lceil 1.5n \rceil,i}(t) + E_{\lfloor 1.5n \rfloor,i}(t)\right] E_{n,i}(t) + \\
& \sum_{ii \neq i}^{m} \chi_{c_{i,i,ii}} E_{n,i}(t) E_{n,ii}(t) \qquad \text{for } 2 < n < k/2, n \text{ odd}
\end{aligned}
\tag{S3}
$$

$$
\begin{aligned}
\frac{dE_{n,i}(t)}{dt} = \mu_i - & \left[\gamma_{a_i} + \delta_{a_i} + \eta_{a_i} + \chi_{a_i} + \phi_{c_i} + \gamma_{c_i} + \delta_{c_i} + \eta_{c_i} + \chi_{c_i} + \mu_i\right] E_{n,i}(t) + \\
& \gamma_{a_i} E_{n+1,i}(t) + \delta_{a_i} E_{n-1,i}(t) + \eta_{a_i} E_{1.5n,i}(t) + \sum_{ii \neq i}^{m} \chi_{a_{i,ii}} E_{n,ii}(t) + \\
& \phi_{c_i} E_{n,i}(t)^2 + \gamma_{c_i} E_{n+1,i}(t) E_{n,i}(t) + \delta_{c_i} E_{n-1,i}(t) E_{n,i}(t) + \eta_{c_i} E_{1.5n,i}(t) E_{n,i}(t) + \\
& \sum_{ii \neq i}^{m} \chi_{c_{i,i,ii}} E_{n,i}(t) E_{n,ii}(t) \qquad \text{for } k/2 \leq n < 3k/4, n \text{ even}
\end{aligned}
\tag{S4}
$$

$$
\begin{aligned}
\frac{dE_{n,i}(t)}{dt} = \mu_i - & \left[\gamma_{a_i} + \delta_{a_i} + \eta_{a_i} + \chi_{a_i} + \phi_{c_i} + \gamma_{c_i} + \delta_{c_i} + \eta_{a_i} + \chi_{c_i} + \mu_i\right] E_{n,i}(t) + \\
& \gamma_{a_i} E_{n+1,i}(t) + \delta_{a_i} E_{n-1,i}(t) + 0.5\eta_{a_i} \left[E_{\lceil 1.5n \rceil,i}(t) + E_{\lfloor 1.5n \rfloor,i}(t)\right] + \\
& \sum_{ii \neq i}^{m} \chi_{a_{i,ii}} E_{n,ii}(t) + \\
& \phi_{c_i} E_{n,i}(t)^2 + \gamma_{c_i} E_{n+1,i}(t) E_{n,i}(t) + \delta_{c_i} E_{n-1,i}(t) E_{n,i}(t) + \\
& 0.5\eta_{c_i} \left[E_{\lceil 1.5n \rceil,i}(t) + E_{\lfloor 1.5n \rfloor,i}(t)\right] E_{n,i}(t) + \sum_{ii \neq i}^{m} \chi_{c_{i,i,ii}} E_{n,i}(t) E_{n,ii}(t) \qquad \text{for } k/2 \leq n < 3k/4, n \text{ odd}
\end{aligned}
\tag{S5}
$$

$$
\begin{aligned}
\frac{dE_{n,i}(t)}{dt} = \mu_i - & \left[\gamma_{a_i} + \delta_{a_i} + \chi_{a_i} + \phi_{c_i} + \gamma_{c_i} + \delta_{c_i} + \chi_{c_i} + \mu_i\right] E_{n,i}(t) + \\
& \gamma_{a_i} E_{n+1,i}(t) + \delta_{a_i} E_{n-1,i}(t) + \sum_{ii \neq i}^{m} \chi_{a_{i,ii}} E_{n,ii}(t) + \\
& \phi_{c_i} E_{n,i}(t)^2 + \gamma_{c_i} E_{n+1,i}(t) E_{n,i}(t) + \delta_{c_i} E_{n-1,i}(t) E_{n,i}(t) + \sum_{ii \neq i}^{m} \chi_{c_{i,i,ii}} E_{n,i}(t) E_{n,ii}(t) \qquad \text{for } 3k/4 \leq n < k
\end{aligned}
\tag{S6}
$$

$$
\begin{aligned}
\frac{dE_{n,i}(t)}{dt} = &\ \mu_i - \left[\delta_{a_i} + \chi_{a_i} + \phi_{c_i} + \delta_{c_i} + \chi_{c_i} + \mu_i\right] E_{n,i}(t) + \\
&\ \delta_{a_i} E_{n-1,i}(t) + \sum_{ii \neq i}^{m} \chi_{a_{i,ii}} E_{n,ii}(t) + \\
&\ \phi_{c_i} E_{n,i}(t)^2 + \delta_{c_i} E_{n-1,i}(t) E_{n,i}(t) + \sum_{ii \neq i}^{m} \chi_{c_{i,i,ii}} E_{n,i}(t) E_{n,ii}(t) \qquad \text{for } n = k
\end{aligned} \tag{S7}
$$

#### S2.2 Lineage probabilities

As with extinction probabilities, we derive SDEs for lineage probabilities (the probability of the subtree descending from lineage $l$ at time $t$, given the lineage is in state $n, i$) using standard notation for SSE models (Maddison et al., 2007). As before, there are different SDEs for boundary cases. The resulting SDEs are:

$$
\begin{aligned}
\frac{dD_{l,n,i}(t)}{dt} = &- \left[\gamma_{a_i} + \rho_{a_i} + \chi_{a_i} + \phi_{c_i} + \gamma_{c_i} + \rho_{a_i} + \chi_{c_i} + \mu_i\right] D_{l,n,i}(t) + \\
&\left[\gamma_{a_i} + \rho_{a_i}\right] D_{l,n+1,i}(t) + \sum_{ii \neq i}^{m} \chi_{a_{i,ii}} D_{l,n,ii}(t) + \\
&\phi_{c_i} D_{l,n,i}(t) E_{n,i}(t) + \left[\gamma_{c_i} + \rho_{c_i}\right] \left[D_{l,n+1,i}(t) E_{n,i}(t) + D_{l,n,i} E_{n+1,i}(t)\right] + \\
&\sum_{ii \neq i} \chi_{c_{i,i,ii}} \left[D_{l,n,ii} E_{n,i}(t) + D_{l,n,i} E_{n,ii}(t)\right] \qquad \text{for } n = 1
\end{aligned} \tag{S8}
$$

$$
\begin{aligned}
\frac{dD_{l,n,i}(t)}{dt} = &- \left[\gamma_{a_i} + \delta_{a_i} + \rho_{a_i} + \eta_{a_i} + \chi_{a_i} + \phi_{c_i} + \gamma_{c_i} + \rho_{c_i} + \eta_{c_i} + \chi_{c_i} + \mu_i\right] D_{l,n,i}(t) + \\
&\gamma_{a_i} D_{l,n+1,i}(t) + \delta_{a_i} D_{l,n-1,i}(t) + \rho_{a_i} D_{l,2n,i}(t) + \eta_{a_i} D_{l,1.5n,i}(t) + \sum_{ii \neq i}^{m} \chi_{a_{i,ii}} D_{l,n,ii}(t) + \\
&\phi_{c_i} D_{l,n,i}(t) E_{n,i}(t) + \\
&\gamma_{c_i} \left[D_{l,n+1,i}(t) E_{n,i}(t) + D_{l,n,i} E_{n+1,i}(t)\right] + \\
&\delta_{c_i} \left[D_{l,n-1,i}(t) E_{n,i}(t) + D_{l,n,i} E_{n-1,i}(t)\right] + \\
&\rho_{c_i} \left[D_{l,2n,i}(t) E_{n,i}(t) + D_{l,n,i} E_{2n,i}(t)\right] + \\
&0.5\eta_{c_i} \left[D_{l,1.5n,i}(t) E_{n,i}(t) + D_{l,n,i} E_{1.5n,i}(t)\right] + \\
&\sum_{ii \neq i}^{m} \chi_{c_{i,i,ii}} \left[D_{l,n,ii} E_{n,i}(t) + D_{l,n,i} E_{n,ii}(t)\right] \qquad \text{for } 2 < n < k/2, n \text{ even}
\end{aligned} \tag{S9}
$$

$$
\begin{aligned}
\frac{dD_{l,n,i}(t)}{dt} = &-\left[\gamma_{a_i} + \delta_{a_i} + \rho_{a_i} + \eta_{a_i} + \chi_{a_i} + \phi_{c_i} + \gamma_{c_i} + \rho_{c_i} + \eta_{c_i} + \chi_{c_i} + \mu_i\right] D_{l,n,i}(t) + \\
&\gamma_{a_i} D_{l,n+1,i}(t) + \delta_{a_i} D_{l,n-1,i}(t) + \rho_{a_i} D_{l,2n,i}(t) + \\
&0.5\eta_{a_i}\left[D_{l,\lceil 1.5n\rceil,i}(t) + D_{l,\lfloor 1.5n\rfloor,i}(t)\right] + \sum_{ii\neq i}^{m} \chi_{a_{i,ii}} D_{l,n,ii}(t) + \\
&\phi_{c_i} D_{l,n,i}(t) E_{n,i}(t) + \\
&\gamma_{c_i}\left[D_{l,n+1,i}(t) E_{n,i}(t) + D_{l,n,i} E_{n+1,i}(t)\right] + \\
&\delta_{c_i}\left[D_{l,n-1,i}(t) E_{n,i}(t) + D_{l,n,i} E_{n-1,i}(t)\right] + \\
&\rho_{c_i}\left[D_{l,2n,i}(t) E_{n,i}(t) + D_{l,n,i} E_{2n,i}(t)\right] + \\
&0.5\eta_{c_i}\left[D_{l,\lceil 1.5n\rceil,i}(t) E_{n,i}(t) + D_{l,n,i} E_{\lceil 1.5n\rceil,i}(t) + D_{l,\lfloor 1.5n\rfloor,i}(t) E_{n,i}(t) + D_{l,n,i} E_{\lfloor 1.5n\rfloor,i}(t)\right] + \\
&\sum_{ii\neq i}^{m} \chi_{c_{i,i,ii}}\left[D_{l,n,ii} E_{n,i}(t) + D_{l,n,i} E_{n,ii}(t)\right] \qquad \text{for } 2 < n < k/2, n \text{ odd}
\end{aligned} \tag{S10}
$$

$$
\begin{aligned}
\frac{dD_{l,n,i}(t)}{dt} = &-\left[\gamma_{a_i} + \delta_{a_i} + \eta_{a_i} + \chi_{a_i} + \phi_{c_i} + \gamma_{c_i} + \eta_{c_i} + \chi_{c_i} + \mu_i\right] D_{l,n,i}(t) + \\
&\gamma_{a_i} D_{l,n+1,i}(t) + \delta_{a_i} D_{l,n-1,i}(t) + \eta_{a_i} D_{l,1.5n,i}(t) + \sum_{ii\neq i}^{m} \chi_{a_{i,ii}} D_{l,n,ii}(t) + \\
&\phi_{c_i} D_{l,n,i}(t) E_{n,i}(t) + \\
&\gamma_{c_i}\left[D_{l,n+1,i}(t) E_{n,i}(t) + D_{l,n,i} E_{n+1,i}(t)\right] + \\
&\delta_{c_i}\left[D_{l,n-1,i}(t) E_{n,i}(t) + D_{l,n,i} E_{n-1,i}(t)\right] + \\
&0.5\eta_{c_i}\left[D_{l,1.5n,i}(t) E_{n,i}(t) + D_{l,n,i} E_{1.5n,i}(t)\right] + \\
&\sum_{ii\neq i}^{m} \chi_{c_{i,i,ii}}\left[D_{l,n,ii} E_{n,i}(t) + D_{l,n,i} E_{n,ii}(t)\right] \qquad \text{for } k/2 < n < 3k/4, n \text{ even}
\end{aligned} \tag{S11}
$$

$$
\begin{aligned}
\frac{dD_{l,n,i}(t)}{dt} = &-\left[\gamma_{a_i} + \delta_{a_i} + \eta_{a_i} + \chi_{a_i} + \phi_{c_i} + \gamma_{c_i} + \eta_{c_i} + \chi_{c_i} + \mu_i\right] D_{l,n,i}(t) + \\
&\gamma_{a_i} D_{l,n+1,i}(t) + \delta_{a_i} D_{l,n-1,i}(t) \\
&0.5\eta_{a_i}\left[D_{l,\lceil 1.5n\rceil,i}(t) + D_{l,\lfloor 1.5n\rfloor,i}(t)\right] + \sum_{ii\neq i}^{m} \chi_{a_{i,ii}} D_{l,n,ii}(t) + \\
&\phi_{c_i} D_{l,n,i}(t) E_{n,i}(t) + \\
&\gamma_{c_i}\left[D_{l,n+1,i}(t) E_{n,i}(t) + D_{l,n,i} E_{n+1,i}(t)\right] + \\
&\delta_{c_i}\left[D_{l,n-1,i}(t) E_{n,i}(t) + D_{l,n,i} E_{n-1,i}(t)\right] + \\
&0.5\eta_{c_i}\left[D_{l,\lceil 1.5n\rceil,i}(t) E_{n,i}(t) + D_{l,n,i} E_{\lceil 1.5n\rceil,i}(t) + D_{l,\lfloor 1.5n\rfloor,i}(t) E_{n,i}(t) + D_{l,n,i} E_{\lfloor 1.5n\rfloor,i}(t)\right] + \\
&\sum_{ii\neq i}^{m} \chi_{c_{i,i,ii}}\left[D_{l,n,ii} E_{n,i}(t) + D_{l,n,i} E_{n,ii}(t)\right] \qquad \text{for } k/2 < n < 3k/4, n \text{ odd}
\end{aligned} \tag{S12}
$$

$$
\begin{aligned}
\frac{dD_{l,n,i}(t)}{dt} = &-\left[\gamma_{a_i} + \delta_{a_i} + \chi_{a_i} + \phi_{c_i} + \gamma_{c_i} + \chi_{c_i} + \mu_i\right] D_{l,n,i}(t) + \\
&\gamma_{a_i} D_{l,n+1,i}(t) + \delta_{a_i} D_{l,n-1,i}(t) + \sum_{ii \neq i}^{m} \chi_{a_{i,ii}} D_{l,n,ii}(t) + \\
&\phi_{c_i} D_{l,n,i}(t) E_{n,i}(t) + \\
&\gamma_{c_i}\left[D_{l,n+1,i}(t) E_{n,i}(t) + D_{l,n,i} E_{n+1,i}(t)\right] + \\
&\delta_{c_i}\left[D_{l,n-1,i}(t) E_{n,i}(t) + D_{l,n,i} E_{n-1,i}(t)\right] + \\
&\sum_{ii \neq i}^{m} \chi_{c_{i,i,ii}}\left[D_{l,n,ii} E_{n,i}(t) + D_{l,n,i} E_{n,ii}(t)\right] \qquad \text{for } 3k/4 < n < k
\end{aligned} \tag{S13}
$$

$$
\begin{aligned}
\frac{dD_{l,n,i}(t)}{dt} = &-\left[\delta_{a_i} + \chi_{a_i} + \phi_{c_i} + + \chi_{c_i} + \mu_i\right] D_{l,n,i}(t) + \\
&\delta_{a_i} D_{l,n-1,i}(t) + \sum_{ii \neq i}^{m} \chi_{a_{i,ii}} D_{l,n,ii}(t) + \\
&\phi_{c_i} D_{l,n,i}(t) E_{n,i}(t) + \\
&\delta_{c_i}\left[D_{l,n-1,i}(t) E_{n,i}(t) + D_{l,n,i} E_{n-1,i}(t)\right] + \\
&\sum_{ii \neq i}^{m} \chi_{c_{i,i,ii}}\left[D_{l,n,ii} E_{n,i}(t) + D_{l,n,i} E_{n,ii}(t)\right] \qquad \text{for } n = k
\end{aligned} \tag{S14}
$$

### S3 Simulation Study

To validate our model, we performed a simulation study where we simulated datasets under biologically-plausible parameter values using our new ChromoHiSSE model and then analyzed those datasets using both ChromoHiSSE—which accommodates process variation—and ChromoSSE—which does not. Our primary goals were (1) to confirm that our model is able to recover the true parameter values and (2) to determine the effect of failing to model process variation (not include hidden states) when process variation is a known generative component of the dataset.

#### S3.1 Simulations

We simulated under two groups of datasets. In the first group of datasets, we kept extinction quite high (we will refer to this as the high-extinction scenario elsewhere in the manuscript), consistent with our empirical estimates. In the second group of datasets we used a lower value for extinction (low-extinction scenario) to test if our ability to recover the true simulating values is affected by high extinction rates.

We performed simulations in `R` using an `R` wrapper for a custom `C++` simulator for the ChromoHiSSE model. For each of the two scenarios, we simulated 100 datasets using the parameter values specified in Table S1. We selected these values based on the estimates from our empirical analysis to ensure that we validated our model under the circumstances most relevant to our results. In some cases, we adjusted the simulating values to simulate datasets that were computationally feasible to analyze at scale. For example, we limited the maximum number of chromosomes in the analysis to 16 (vs. 72 in the empirical analysis). This significantly reduced the state space (and thus runtime) of the ChromoHiSSE model used to analyze those datasets. Because we lowered the maximum chromosome number, we also lowered the higher set of anagenetic and cladogenetic dysploidy rates (in hidden state *i*). We retained simulations that contained at least 10% of tips in the lower-represented hidden state and that had between 450 and 550 tips. We then analyzed those datasets using ChromoHiSSE and ChromoSSE in `RevBayes`. The simulating code, simulated datasets, and analysis code is available at Zenodo DOI: 10.5281/zenodo.8320249.

**Table S1:** True parameter values used to simulated datasets for the high- and low-extinction scenarios. Parameters that vary across the two scenarios are in bold.

| parameter | high extinction | low extinction |
|---|---|---|
| **Time** | **25** | **10** |
| Max chromo | 16 | 16 |
| Num hidden | 2 | 2 |
| $\gamma_{a_i}$ | 4.6 | 4.6 |
| $\delta_{a_i}$ | 4.2 | 4.2 |
| $\gamma_{a_{ii}}$ | 0.15 | 0.15 |
| $\delta_{a_{ii}}$ | 0.12 | 0.12 |
| $\chi_a$ | 0.02 | 0.02 |
| $\phi_{c_i}$ | 0.07 | 0.07 |
| $\delta_{c_{ii}}$ | 0.5 | 0.5 |
| $\gamma_{c_i}$ | 0.3 | 0.3 |
| $\phi_{c_{ii}}$ | 0.9 | 0.9 |
| $\delta_{c_{ii}}$ | 0.08 | 0.08 |
| $\gamma_{c_{ii}}$ | 0.07 | 0.07 |
| $\chi_c$ | 0.1 | 0.1 |
| $\mu$ | **0.9** | **0.3** |

#### S3.2 Results

##### S3.2.1 High extinction rates

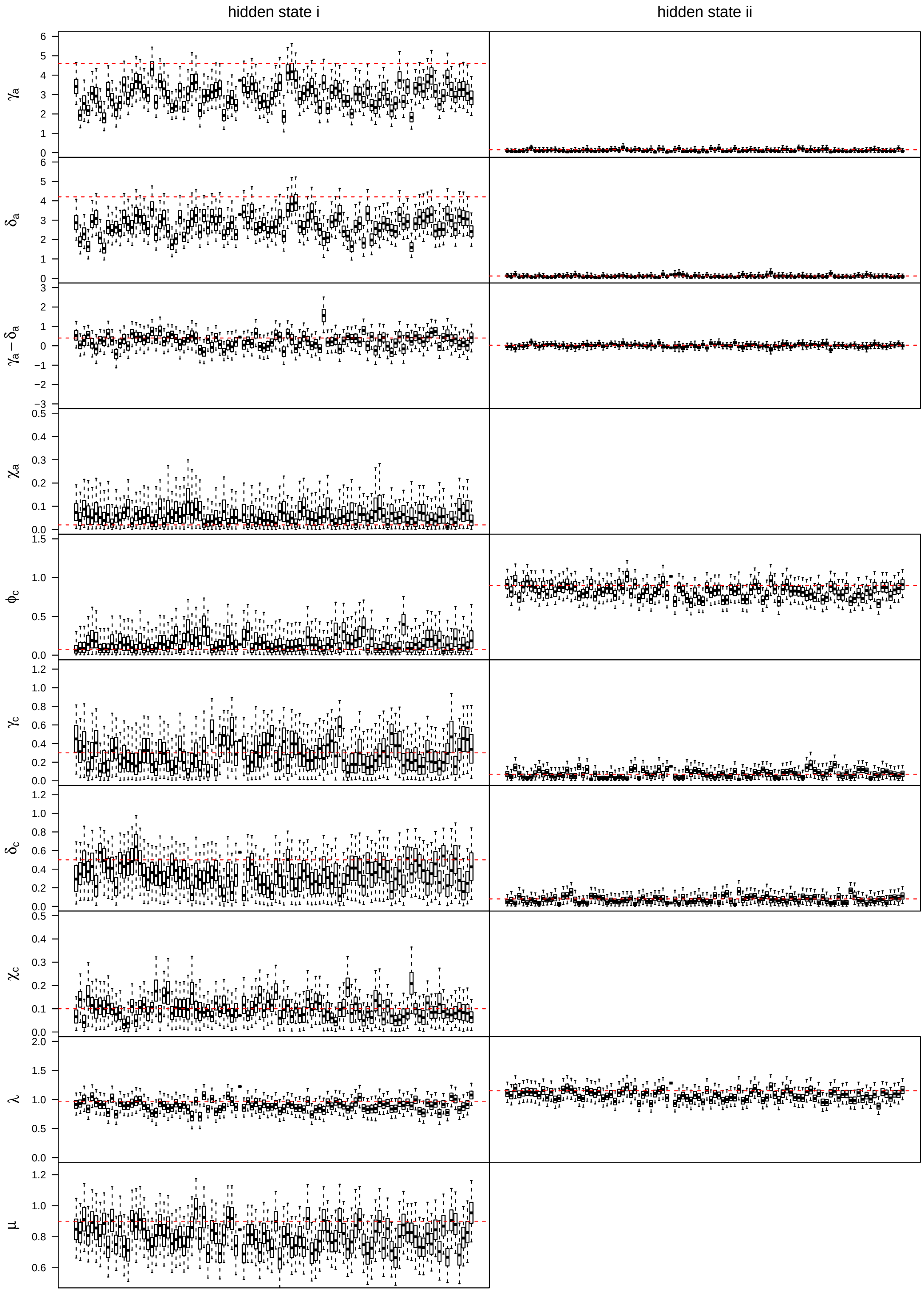


**Figure S1: Absolute parameter estimates under the true model.** For each parameter (in rows), we plot posterior distributions for each simulation (boxplots) for both hidden states *i* and *ii* (columns, when relevant to the parameter). For each boxplot, the whiskers represent the 95% HPD, boxes represent the 50% HPD, and black bars represent the posterior median. Dashed red lines correspond the true values.

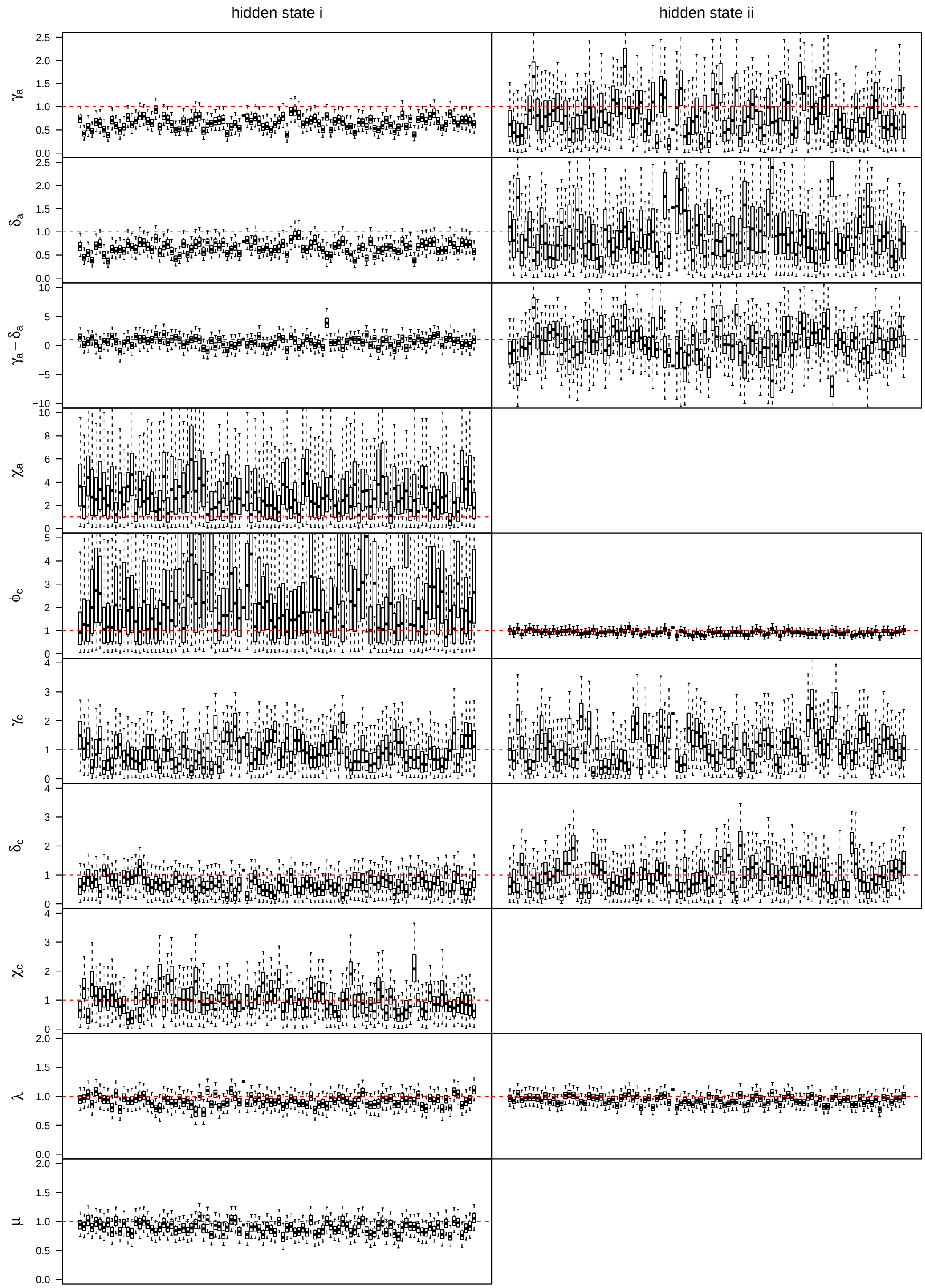


**Figure S2: Relative parameter estimates under the true model.** For each parameter (in rows), we plot posterior distributions for each simulation (boxplots) for both hidden states *i* and *ii* (columns, when relevant to the parameter). For each boxplot, the whiskers represent the 95% HPD, boxes represent the 50% HPD, and black bars represent the posterior median. Sampled parameter values are normalized by dividing by the true parameter value, so these boxplots are relative to the true value. Dashed red lines are at 1, representing the true value divided by itself.

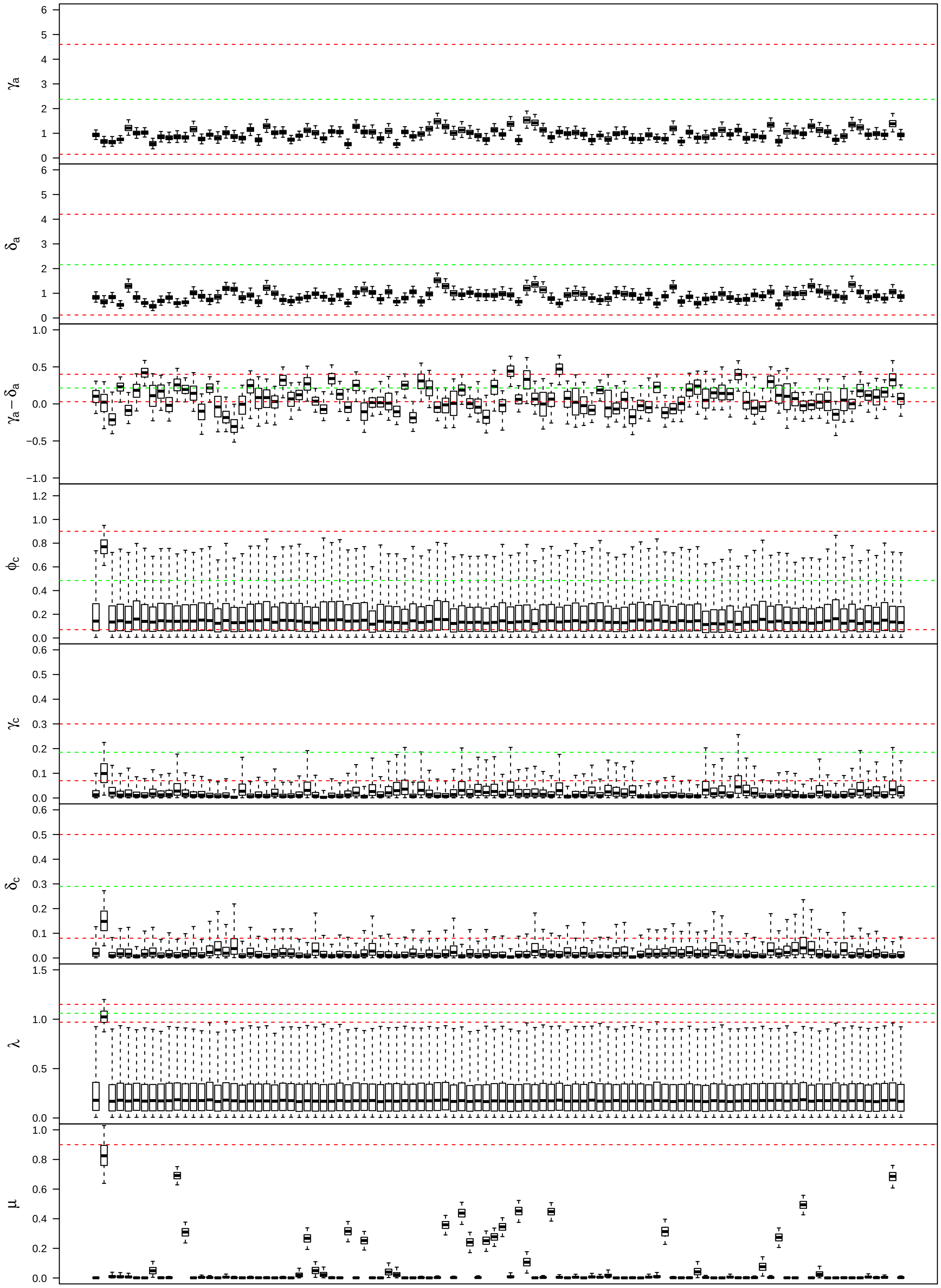


**Figure S3: Absolute parameter estimates under the ChromoSSE model (*i.e.*, no hidden states).** For each parameter in the ChromoSSE model, we plot posterior distributions for each simulation (boxplots). For each boxplot, the whiskers represent the 95% HPD, boxes represent the 50% HPD, and black bars represent the posterior median. Dashed red lines correspond to the true values for the two hidden states; dashed green lines are the average of the true values.

**Table S2:** Parameter estimates, error, and coverage (how often the true value falls within the 95% confidence interval) under the ChromoHiSSE model.

| parameter | percent error | squared error (relative) | coverage |
|---|---|---|---|
| $\gamma_{ai}$ | 34.46 | 0.61 | 0.26 |
| $\delta_{ai}$ | 34.26 | 0.55 | 0.32 |
| $\gamma_{ai} - \delta_{ai}$ | 63.80 | 0.28 | 0.94 |
| $\gamma_{aii}$ | 31.89 | 0.02 | 0.95 |
| $\delta_{aii}$ | 30.10 | 0.02 | 0.98 |
| $\gamma_{aii} - \delta_{aii}$ | 188.11 | 0.18 | 0.97 |
| $\chi_a$ | 208.43 | 0.11 | 0.99 |
| $\phi_{ci}$ | 136.08 | 0.20 | 0.99 |
| $\gamma_{ci}$ | 29.71 | 0.04 | 0.99 |
| $\delta_{ci}$ | 32.11 | 0.07 | 0.94 |
| $\phi_{cii}$ | 9.69 | 0.01 | 0.88 |
| $\gamma_{cii}$ | 39.46 | 0.02 | 0.96 |
| $\delta_{cii}$ | 29.92 | 0.01 | 0.97 |
| $\chi_c$ | 25.67 | 0.01 | 0.99 |
| $\lambda_i$ | 9.55 | 0.01 | 0.93 |
| $\lambda_{ii}$ | 7.30 | 0.01 | 0.90 |
| $\mu$ | 11.08 | 0.02 | 0.89 |

**Table S3:** Parameter estimates, error, and coverage (how often the true value falls within the 95% confidence interval) under the ChromoSSE model (*i.e.*, no hidden states).

| parameter | percent error | squared error (relative) | coverage |
|---|---|---|---|
| $\gamma_a$ | 58.82 | 0.84 | 0.00 |
| $\delta_a$ | 58.25 | 0.75 | 0.00 |
| $\gamma_a - \delta_a$ | 79.95 | 0.19 | 0.74 |
| $\phi_c$ | 58.96 | 0.17 | 0.99 |
| $\gamma_c$ | 85.84 | 0.14 | 0.10 |
| $\delta_c$ | 90.98 | 0.24 | 0.00 |
| $\lambda$ | 75.74 | 0.61 | 0.01 |
| $\mu$ | 91.14 | 0.78 | 0.01 |

##### S3.2.2 Low extinction rates

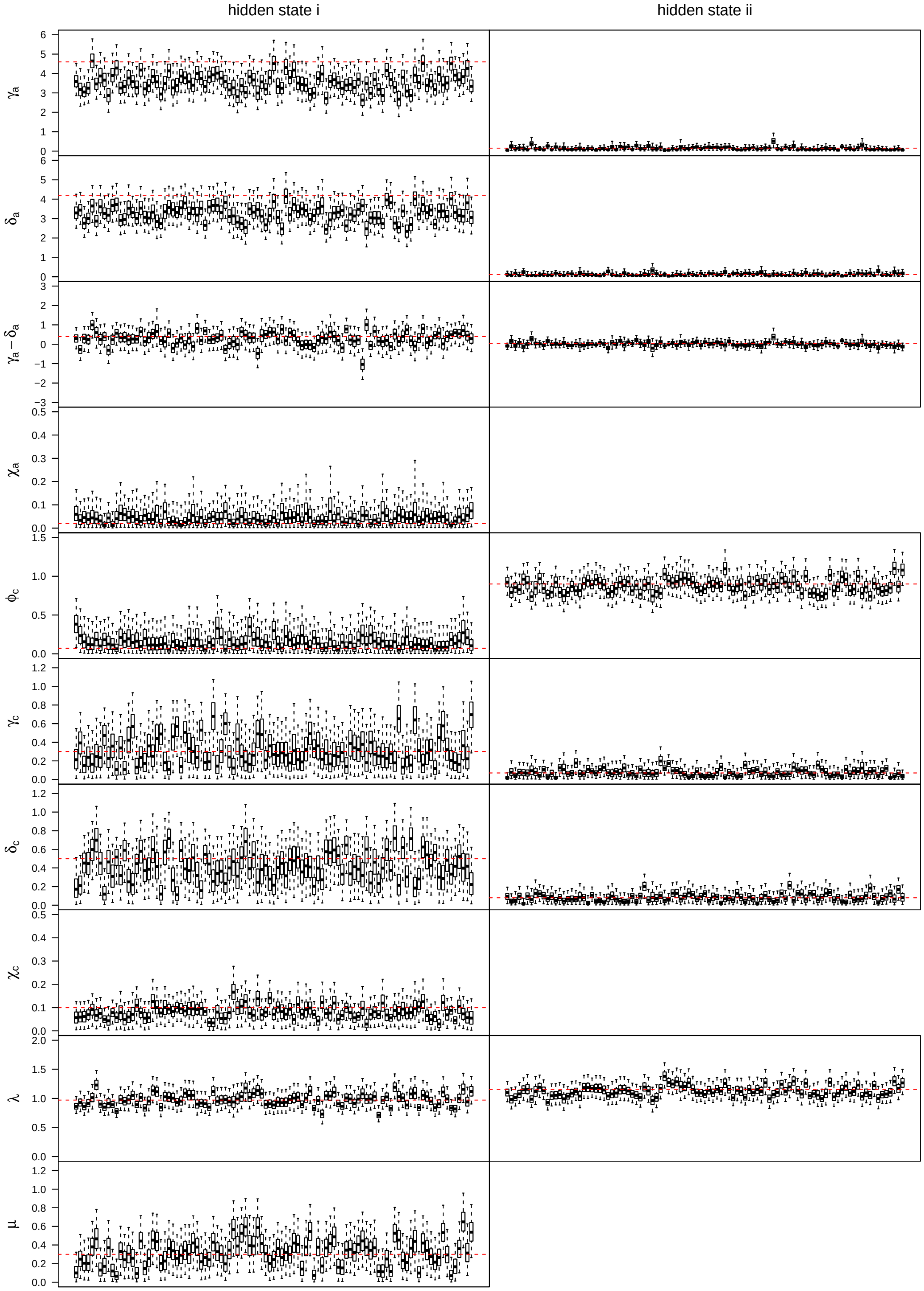


**Figure S4: Absolute parameter estimates under the true model.** For each parameter (in rows), we plot posterior distributions for each simulation (boxplots) for both hidden states *i* and *ii* (columns, when relevant to the parameter). For each boxplot, the whiskers represent the 95% HPD, boxes represent the 50% HPD, and black bars represent the posterior median. Dashed red lines correspond the true values.

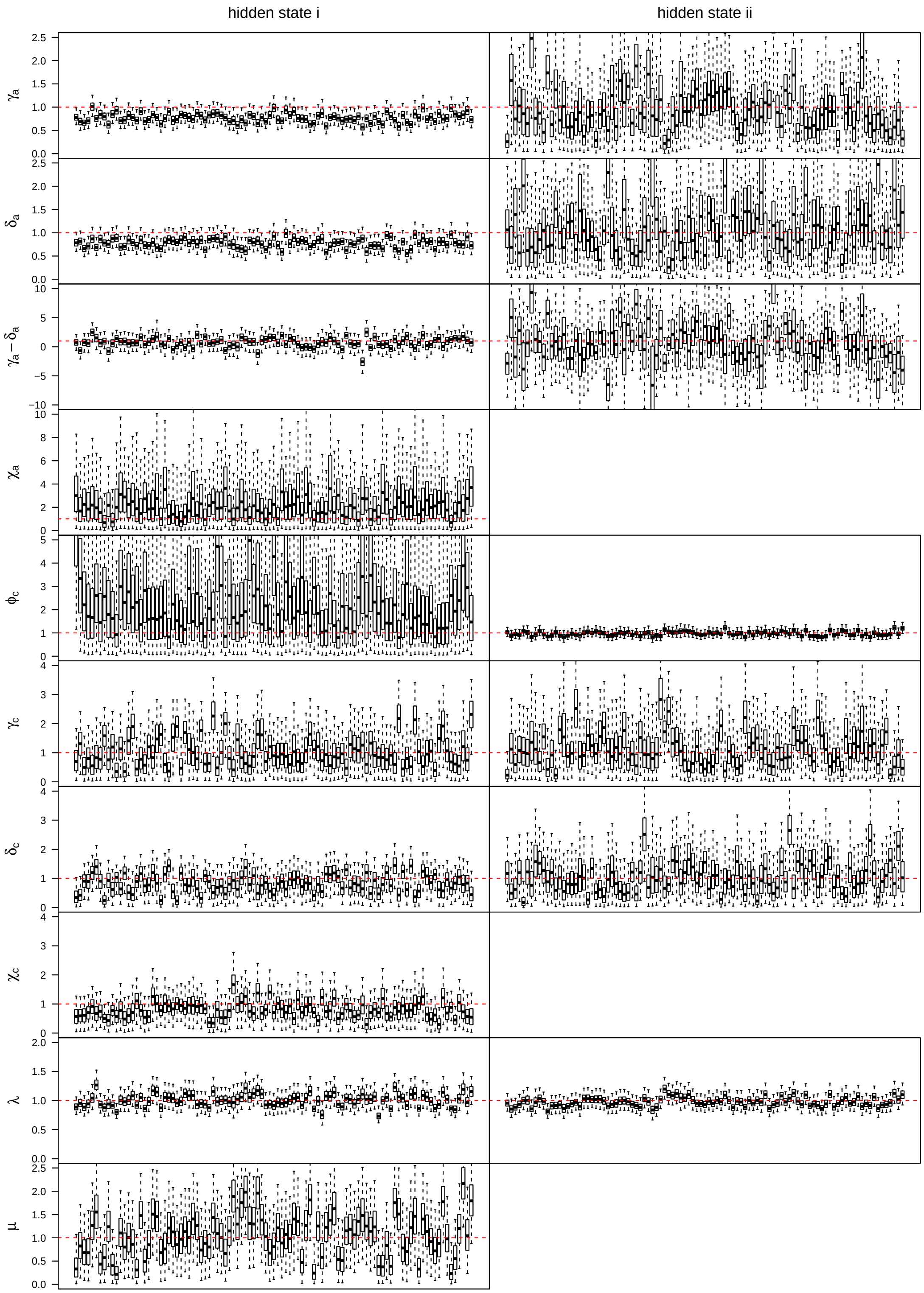


**Figure S5: Relative parameter estimates under the true model.** For each parameter (in rows), we plot posterior distributions for each simulation (boxplots) for both hidden states *i* and *ii* (columns, when relevant to the parameter). For each boxplot, the whiskers represent the 95% HPD, boxes represent the 50% HPD, and black bars represent the posterior median. Sampled parameter values are normalized by dividing by the true parameter value, so these boxplots are relative to the true value. Dashed red lines are at 1, representing the true value divided by itself.

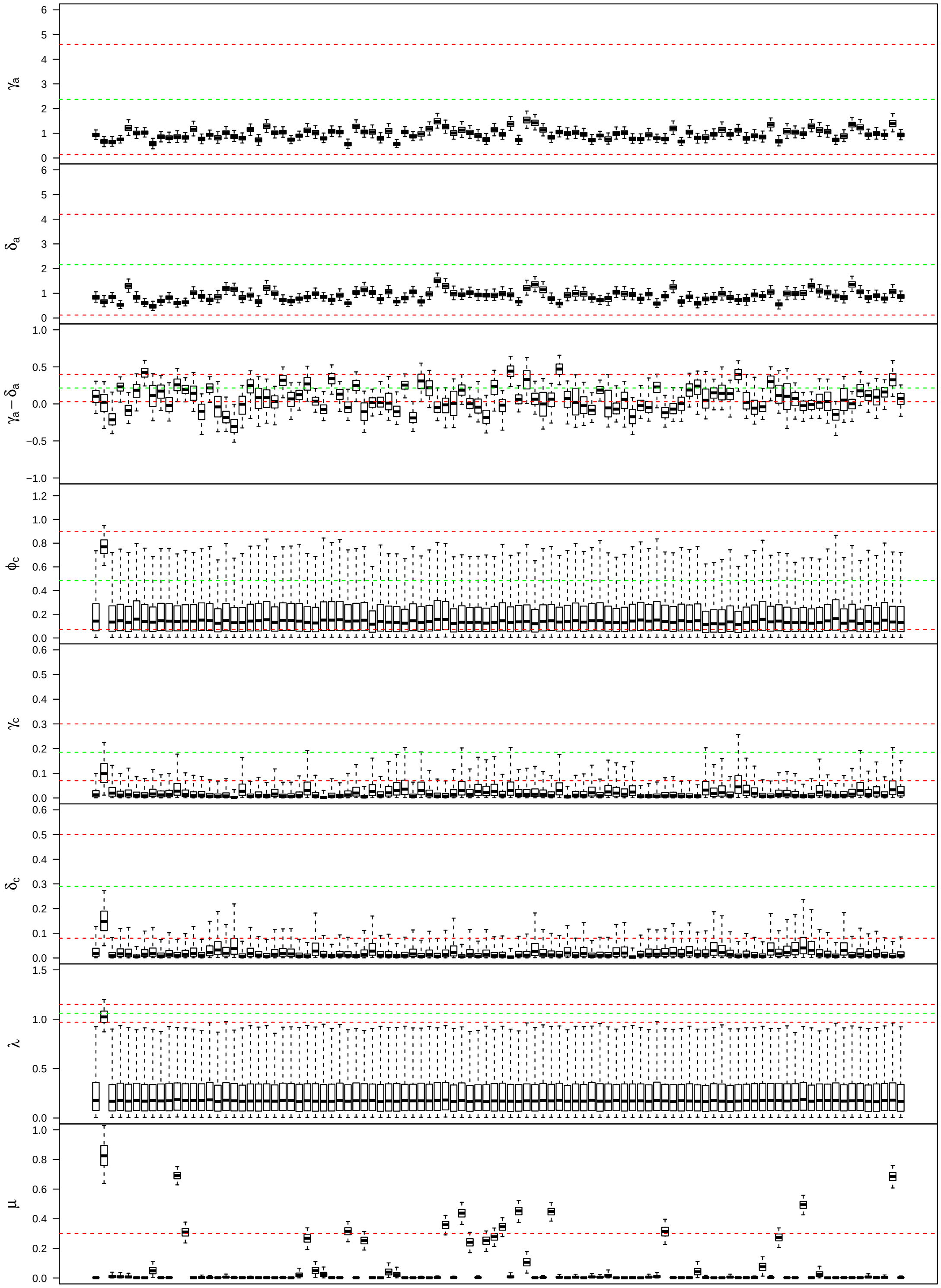


**Figure S6: Absolute parameter estimates under the ChromoSSE model (*i.e.*, no hidden states).** For each parameter in the ChromoSSE model, we plot posterior distributions for each simulation (boxplots). For each boxplot, the whiskers represent the 95% HPD, boxes represent the 50% HPD, and black bars represent the posterior median. Dashed red lines correspond to the true values for the two hidden states; dashed green lines are the average of the true values.

**Table S4:** Parameter estimates, error, and coverage (how often the true value falls within the 95% confidence interval) under the ChromoHiSSE model.

| parameter | percent error | squared error (relative) | coverage |
|---|---|---|---|
| $\gamma_{ai}$ | 22.38 | 0.27 | 0.52 |
| $\delta_{ai}$ | 21.91 | 0.24 | 0.62 |
| $\gamma_{ai} - \delta_{ai}$ | 64.54 | 0.28 | 0.95 |
| $\gamma_{aii}$ | 33.96 | 0.03 | 0.94 |
| $\delta_{aii}$ | 34.97 | 0.03 | 0.99 |
| $\gamma_{aii} - \delta_{aii}$ | 242.23 | 0.30 | 0.99 |
| $\chi_a$ | 138.36 | 0.05 | 1.00 |
| $\phi_{ci}$ | 137.89 | 0.19 | 0.99 |
| $\gamma_{ci}$ | 34.04 | 0.06 | 1.00 |
| $\delta_{ci}$ | 27.02 | 0.05 | 0.95 |
| $\phi_{cii}$ | 7.78 | 0.01 | 1.00 |
| $\gamma_{cii}$ | 38.40 | 0.02 | 0.98 |
| $\delta_{cii}$ | 34.85 | 0.02 | 0.98 |
| $\chi_c$ | 26.19 | 0.01 | 0.93 |
| $\lambda_i$ | 8.30 | 0.01 | 0.95 |
| $\lambda_{ii}$ | 6.34 | 0.01 | 0.98 |
| $\mu$ | 35.07 | 0.06 | 0.94 |

**Table S5:** Parameter estimates, error, and coverage (how often the true value falls within the 95% confidence interval) under the ChromoSSE model (*i.e.*, no hidden states).

| parameter | percent error | squared error (relative) | coverage |
|---:|---:|---:|---:|
| $\gamma_a$ | 28.41 | 0.27 | 0.34 |
| $\delta_a$ | 29.78 | 0.28 | 0.37 |
| $\gamma_a - \delta_a$ | 176.23 | 1.00 | 0.72 |
| $\phi_c$ | 19.07 | 0.04 | 0.86 |
| $\gamma_c$ | 58.04 | 0.07 | 0.89 |
| $\delta_c$ | 64.35 | 0.13 | 0.72 |
| $\lambda$ | 33.85 | 0.14 | 0.99 |
| $\mu$ | 82.29 | 0.23 | 0.18 |

#### S4 ChromoSSE Results (no hidden state)

In addition to the analyses presented in the manuscript, we ran four additional ChromoSSE analyses to test the effects of including or excluding polyploidy and including or excluding the *Siderostictae* clade. *Siderostictae* is sister to the rest of *Carex* and has documented polyploidy. We thus wanted to know if we had the power to model polyploidy when including *Siderostictae* and whether including polyploidy, with or without *Siderostictae*, was feasible given the low rates documented in the rest of *Carex*. It was not feasible to run these tests using ChromoHiSSE given the computational limits of the analysis, so we used ChromoSSE instead.

##### S4.1 With *Carex* subg. *Siderostictae*, with polyploidy

When we include *Siderostictae* and $\rho_a$, the estimate for $\rho_a$ approximates 0. It thus appears that we do not have the power to meaningfully detect polyploidy even when including the clade that contains known polyploid lineages.

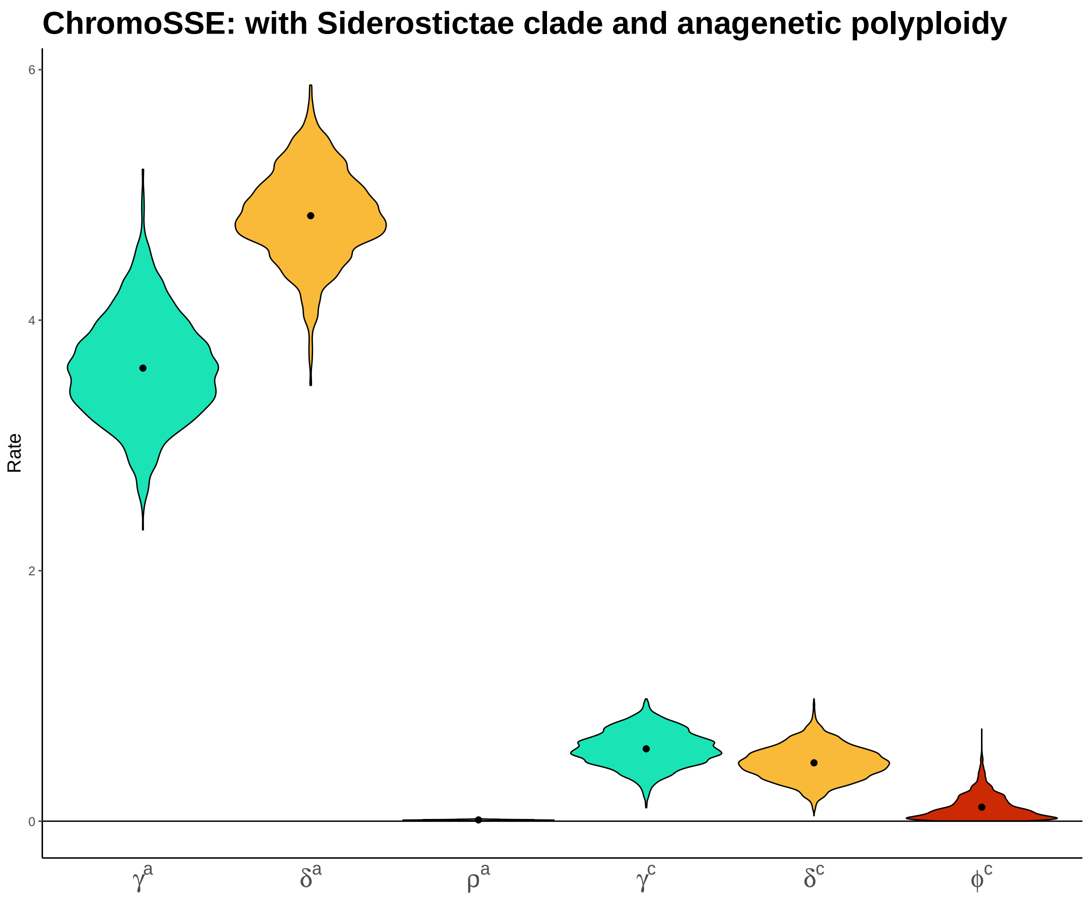


**Figure S7:** Posterior distributions of rate estimates from a ChromoSSE model that allows anagenetic polyploidy ($\rho_a$; estimated to be ∼0. Polyploidy events are rare for *Carex*.) on data that includes *Carex* subg. *Siderostictae*.

#### S4.2 With *Carex* subg. *Siderostictae*, without polyploidy

When we include *Siderostictae* but do not include polyploidy, we recover similar parameter estimates as in the other ChromoSSE analyses. It thus does not appear to *hurt* the analysis to include rare polyploidy events without directly modeling polyploidy, but as it is technically incorrect, we decided to drop *Siderostictae* from the main analysis using ChromoHiSSE.

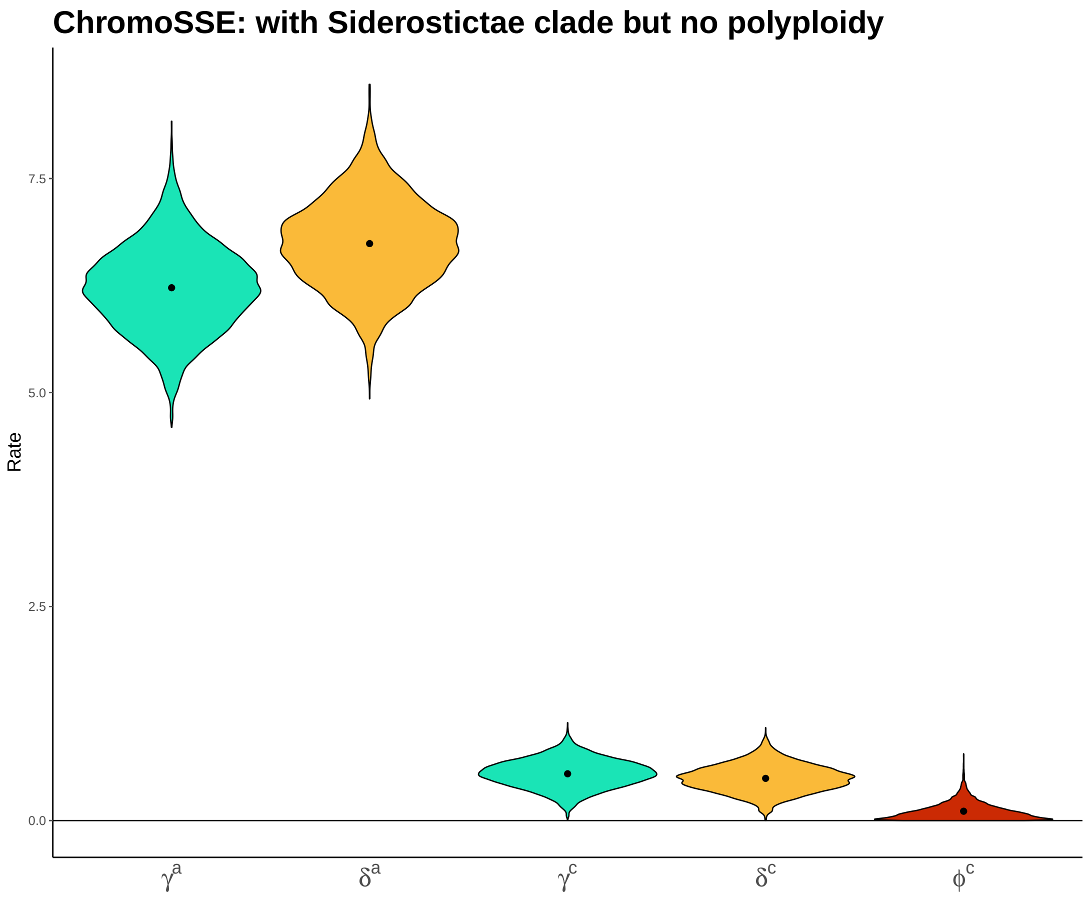


**Figure S8:** Posterior distributions of rate estimates from a ChromoSSE model that does not allow anagenetic polyploidy on data that includes *Carex* subg. *Siderostictae*. Results show that cladogenesis with or without chromosome numbers happens at the same pace since posterior distributions of cladogenetic rates for fusions, fissions, and no character change overlap.

#### S4.3 Without *Carex* subg. *Siderostictae*, with polyploidy

Similar to S4.1, our estimate for $\rho_a$ approximates 0. Including a polyploidy parameter ($\rho_a$) does not seem to meaningfully affect the estimates of other parameters.

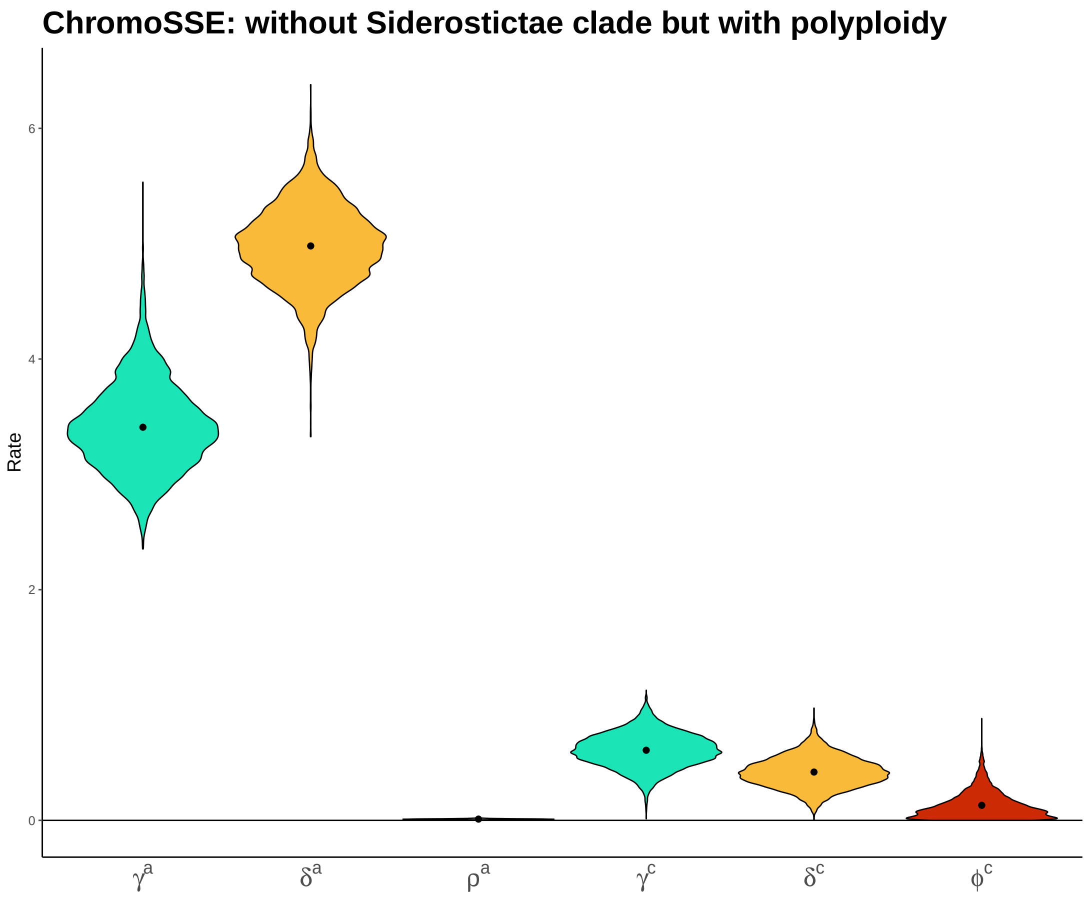


**Figure S9:** Posterior distributions of rate estimates from a ChromoSSE model that allows anagenetic polyploidy ($\rho_a$; estimated to be ∼0. Polyploidy events are rare for *Carex*.) on data that does not include *Carex* subg. *Siderostictae*.

##### S4.4 Without *Carex* subg. *Siderostictae*, without polyploidy

Finally, we tested excluding both $\rho_a$ (polyploidy) and the *Siderostictae* clade. This ChromoSSE analysis most closely resembles the scenarios for our ChromoHiSSE analysis presented in the main text of the manuscript.

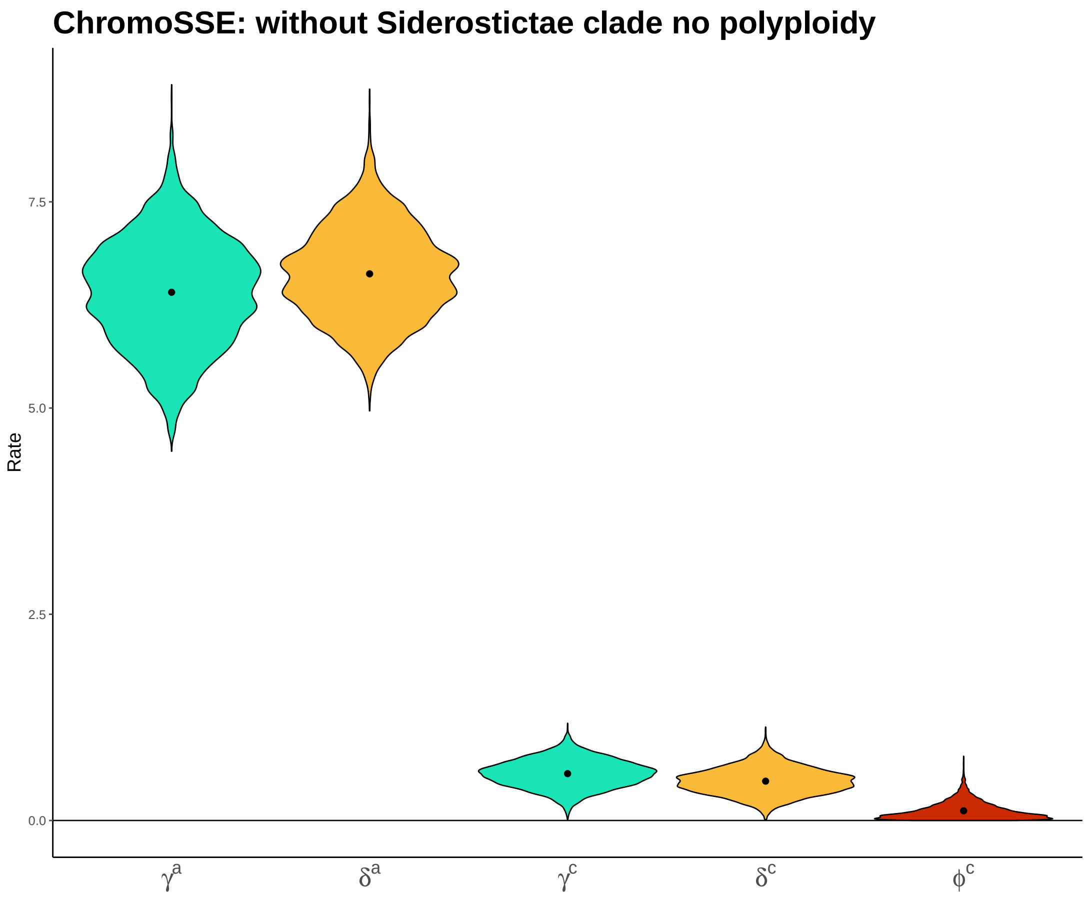


**Figure S10:** Posterior distributions of rate estimates from a ChromoSSE model that does not allow anagenetic polyploidy on data that does not include *Carex* subg. *Siderostictae*. Overlapping parameters of cladogenesis.
